# Telomere-to-telomere assembly and haplotype analysis of tetraploid *Dendrobium officinale* illuminate Orchidaceae polyploid evolution and mycorrhizal symbiosis genes

**DOI:** 10.64898/2026.03.04.709700

**Authors:** Enlian Chen, Jialu Xu, Yuansheng Liu, Yongliang Li, Yiping Feng, Qinghua Lu, Xiaoyu Ding, Zhitao Niu, Si Qin, Shance Niu, Yibo Luo, Xinhong Guo, Xiao Luo

## Abstract

*Dendrobium officinale* is a typical epiphytic orchid. We report the telomere-to-telomere (T2T) genome assembly for *D. officinale*, representing the first T2T reference genome within the Orchidaceae family. The assembly is anchored to 19 chromosomes and contains 38 complete telomeres and 15 characterized centromeres. We further generated haplotype-resolved assemblies of the autotetraploid genome, identifying 12,761 sets of tetra-allelic genes. Based on synonymous substitution analysis, we inferred that the autotetraploidization event occurred approximately 0.86 million years ago. A systematic analysis of the SWEET gene family across the genus *Dendrobium* revealed that the gene family size is shaped primarily by epiphytic types and environmental factors. In *D. officinale* from Langshan, eight *SWEET* genes were specifically expressed in roots, suggesting they may play specialized roles in the root mycorrhizal system, potentially contributing to the *D. officinale*’s ability to recruit and maintain fungal partners. Together, these resources provide valuable foundations for studies of orchid evolution, functional genomics, and molecular breeding.

## Introduction

*Dendrobium*, one of the largest genera in the Orchidaceae family, comprises approximately 1,500 species mainly distributed across Asia and Oceania(1; 2). Among them, *Dendrobium officinale* Kimura et Migo, a perennial herbaceous species, is rich in various bioactive compounds and exhibits diverse pharmacological activities, including antioxidant, hypoglycemic, hepatoprotective, anti-atherosclerotic, and osteoporosis-preventive effects, earning it the reputation of the “immortal herb”(3; 4; 5; 6; 7). Nevertheless, wild *D. officinale* predominantly grows epiphytically on cliff rocks or tree trunks under extremely harsh environmental conditions, resulting in limited natural populations and rendering it a highly scarce natural resource(8; 9).

A high-quality reference genome of *D. officinale* is essential for downstream biological research and breeding, enabling insights into the genetic mechanisms underlying bioactive traits and facilitating targeted trait improvement. To date, three versions of the diploid *D. officinale* genome assembly have been published, each representing incremental improvements but still with notable limitations. The initial assembly, released in 2015, was generated from plant material collected in Puer, Yunnan province, and resulted in a genome size of 1.35 Gbp (10). In 2016, an alternative assembly was constructed from material collected in Guangnan county, Yunnan province, yielding a smaller 1.01 Gbp genome with improved contiguity (11). The first chromosome-level assembly, published in 2021, was based on material collected in Huoshan county, Anhui province, and generated a 1.23 Gbp genome with significantly enhanced structural continuity. Nevertheless, this assembly still consisted of 2,430 contigs, highlighting persistent challenges in fully resolving complex or repetitive genomic regions (12). Although these genomic resources have facilitated preliminary investigations into genes underlying medicinal compound biosynthesis, the absence of a highly contiguous, high-quality reference genome has constrained comprehensive characterization of pharmacologically active biosynthetic pathways and limited advanced genomic research critical for improving *D. officinale* cultivation. Moreover, current genomic studies have primarily focused on diploid *D. officinale* (2n = 38), whereas investigations on polyploid counterparts remain scarce. To date, no genome sequence has been reported for wild tetraploid *D. officinale*, and existing studies have largely relied on colchicine-induced polyploid lines, focusing on physiological traits and metabolite accumulation comparisons between diploid and tetraploid plants (13; 14). Consequently, the lack of high-quality genomic resources for tetraploid *D. officinale* continues to hinder deeper insights into its genome architecture and potential applications in breeding.

Like many other orchids, the growth and development of *D. officinale* strongly depend on symbiotic associations with orchid mycorrhizal fungi (OMF). During seed germination, orchid seeds lack an endosperm and therefore require fungi to provide carbon sources to support seed germination and subsequent protocorm development(15; 16). Previous studies have reported that orchids mainly obtain carbon from fungi in the form of glucose. This process involves the hydrolysis of fungal trehalose by orchid-specific trehalases to release glucose, which is subsequently transported by sugar transporters, among which members of the SWEET family may play critical roles(17). In mature *D. officinale* plants, roots continue to maintain associations with mycorrhizal fungi, from which the plant obtains nutrients(18). This root–fungus interaction not only provides nutritional support for *D. officinale* but also underpins its capacity to survive and adapt to epiphytic environments.

In this study, we assembled the first telomere-to-telomere, reference-quality genome of tetraploid *D. officinale*, together with four haplotype-resolved chromosome-level assemblies. Using these resources, we performed comprehensive phylogenomic and comparative genomic analyses, and revealed a recent whole-genome duplication (WGD) event. We further expanded our analyses to the SWEET gene family across 19 *Dendrobium* species, investigating gene copy numbers and clade-specific distribution patterns. In *D. officinale*, eight *SWEET* genes were found to be exclusively expressed in roots, suggesting potential involvement in root–fungus associations. Together, these results not only provide novel insights into genome evolution and functional diversification of SWEET transporters in *Dendrobium*, but also establish valuable genomic resources for evolutionary studies and molecular breeding of medicinal orchids.

## Results

### Genome assembly of *D. officinale*

We generated a comprehensive sequencing dataset for *D. officinale* from Langshan Mountain, Hunan Province, comprising 168.4 Gbp of Oxford Nanopore (ONT) long reads, including 68.0 Gbp of ultra-long reads (read N50 = 100 kbp), as well as 167.1 Gbp of PacBio HiFi reads, 226.0 Gbp of Hi-C paired-end reads, and 120.0 Gbp of DNBSEQ-T7 short reads (Supplementary Table 1). Flow cytometry estimated the nuclear DNA content to be approximately 4,404.30 Mbp (Supplementary Figure 1A). K-mer frequency analysis (k=19) using GenomeScope supported an autotetraploid genome architecture and suggested a haploid genome size of 1.01 Gbp (Supplementary Figure 1B).

To construct a reference-grade genome, we performed de novo assembly using hifiasm (19), integrating PacBio HiFi reads and ONT ultra-long reads. After removal of duplicated sequences and polishing, we obtained a highly contiguous primary contig set comprising 130 contigs. These contigs were subsequently anchored and ordered into chromosomes using Hi-C chromatin interaction data. Hi-C reads were aligned to the contigs with BWA (20), and chromosome-scale scaffolding was performed using HapHiC (21), achieving a Hi-C anchoring rate of 99.99%. Manual curation with Juicebox (22) was then used to verify contig order/orientation and to correct potential structural inconsistencies, producing 19 pseudochromosomes (Supplementary Figure 2).

Residual gaps were closed using TGS-GapCloser (23) with both ONT and HiFi reads, resulting in a gap-free collapsed assembly. By searching for the canonical telomeric repeats “TTTAGGG/CCCTAAA”, we identified telomeric repeats at 38 chromosome ends (Figure 1B; Supplementary Table 2). The final telomere-to-telomere (T2T) collapsed assembly spans 1,148,476,025 bp and shows high base-level accuracy and completeness (Table 1).

**Figure 1.**
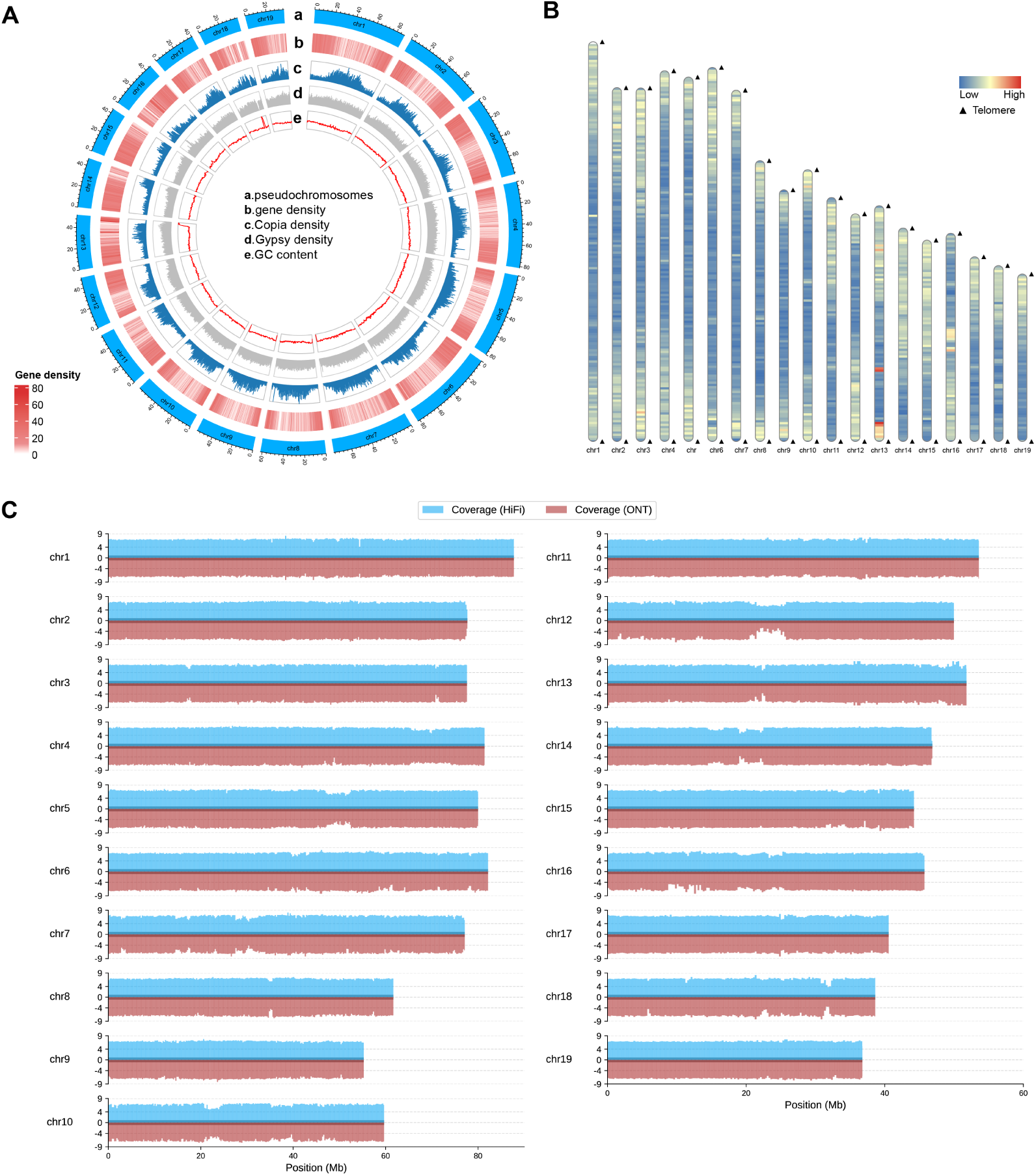
Telomere-to-telomere genome asembly of *Dendrobium officinale*. (A) Genome features, including chromosome pseudomolecule structure (a), gene density (b), Copia density (c), Gypsy density (d), and GC content (e). Densities were calculated using 500-kbp sliding windows. (B) Telomere detection map. Black triangles indicate the positions of telomeres on assembled chromosomes. The color gradient represents gene density. (C) Read coverage of ONT and PacBio HiFi reads on the *D. officinale* T2T genome assembly, with the y-axis representing log_2_-mapping depth.

**Table 1.** Summary of the T2T assembly and haplotype-resolved assemblies of the tetraploid *D. officinale*.

| Assembly | *D. officinale*<br>v3.0<sup>a</sup> | T2T<br>genome | hapA<br>genome | hapB<br>genome | hapC<br>genome | hapD<br>genome |
| --- | --- | --- | --- | --- | --- | --- |
| Number of chromosomes | 19 | 19 | 19 | 19 | 19 | 19 |
| Number of gaps | 1158 | 0 | 2881 | 2734 | 2772 | 2855 |
| Assembly Length (Gbp) | 1.23 | 1.15 | 1.03 | 1.01 | 1.02 | 1.02 |
| Contig N50 (Kbp) | 1,436.1 | 61,670.2 | 575.3 | 606.4 | 596.4 | 586.6 |
| Number of telomeres | - | 38 | - | - | - | - |
| Number of centromeres | - | 15 | - | - | - | - |
| Number of gene models | 27631 | 30718 | 25296 | 25395 | 26110 | 26018 |
| GC Content (%) | 35.5 | 35.34 | 35.04 | 35.03 | 35.04 | 35.03 |
| BUSCOs (%) | 91.2 | 96.8 | 90.8 | 91.0 | 90.3 | 90.8 |
| QV | - | 43.03 | 30.08 | 30.23 | 30.04 | 30.02 |
| LAI | - | 14.26 | 12.18 | 12.23 | 12.27 | 12.13 |
| HiFi reads mapped rate (%) | - | 99.99 | 99.96 | 99.96 | 99.96 | 99.96 |

a: The *D. officinale* v3.0 genome assembly, published by Niu, Zhitao, et al. (2021)(12), is available under the GenBank assembly accession number GCA\_019514585.1. Note that the samples of *D. officinale* v3.0 are from Huoshan, Anhui Province, China.

### Haplotype-resolved assembly

To resolve haplotype-specific variation in this autotetraploid genome, we partitioned HiFi reads into four haplotype-enriched bins based on variant information against the collapsed reference, assembled each bin independently, and anchored haplotype contigs into a consistent 19-chromosome framework using Hi-C data (Supplementary Figure 3). The resulting haplotype assemblies (hapA–hapD) span 1.01–1.03 Gbp, with BUSCO completeness of 90.3–91.0% and QV of 30.02–30.23 (Table 1). These assemblies provide a chromosome-scale haplotype-resolved genomic resource for tetraploid *D. officinale*.

### Evaluation of *D. officinale* genome assembly

Comparative genomics against the *D. officinale* v3.0 reference was performed to assess the quality of the T2T assembly. Whole-genome alignments and gene-based comparisons both exhibited strong synteny along the expected diagonal (Supplementary Figures 4,5), confirming structural fidelity and supporting the continuity of the assembly relative to the previous reference. To further evaluate genome completeness, BUSCO(24) analysis was performed using the embryophyta odb10 dataset. The results showed that 1,563 genes (96.84% of the total conserved genes) were identified in the T2T genome. Among these, 1,509 genes were annotated as complete single-copy genes, while 54 were annotated as complete duplicated genes. The overall query coverage of the HiFi and ONT read alignments was 99.99% and 93.29%, respectively (Figure 1C). Using Merqury(25), the quality value (QV) of the T2T assembly was estimated at 43.03, and the LTR Assembly Index (LAI) was 14.26, both indicating high assembly continuity and completeness.

We comprehensively assessed the assembly quality of four haploid genomes. BUSCO analysis revealed completeness scores ranging from 90.3% to 91.0%, indicating high completeness of conserved single-copy orthologs across all haplotypes. LAI values ranged from 12.13 to 12.27, indicating good assembly continuity and resolution of repetitive regions. Furthermore, QV scores ranged from 30.02 to 30.23, demonstrating high base-level accuracy. All four haplotypes showed a HiFi read mapping rate of 99.96%. Collectively, these results demonstrate that each haplotype assembly exhibits high completeness, continuity, and accuracy, providing a robust foundation for further comparative genomic analyses (Table 1, Supplementary Figure 6). Consistent with these quality assessments, JCVI comparisons between *D. officinale* v3.0 and the four haplotype assemblies revealed strong gene collinearity, supporting the structural consistency of the haplotype genomes (Supplementary Figure 7).

### Genome annotation and gene prediction

Tandem repeat analysis using TRF(26) identified satellite, minisatellite, and microsatellite sequences comprising 2.49% of the genome. Repetitive elements accounted for 74.46% of the assembled genome, with Class I Retrotransposons representing the largest fraction (48.39%), predominantly LTR retrotransposons. Within LTRs, the Gypsy lineage was the most abundant (31.70%), followed by Copia (7.59%). DNA transposons covered 15.76% of the genome (Figure 1A, Supplementary Table 3).

Gene structure annotation of the T2T genome was performed using a combination of homology-based, de novo, and transcriptome-based annotation methods, identifying 30,718 protein-coding genes and 42,410 transcripts. The annotated protein-coding genes had an average gene length of 11,860.08 bp, an average CDS length of 1,090.21 bp, and an average of four exons per gene. To assess annotation completeness, we used a set of 1,614 conserved BUSCO genes from the embryophyta odb10 database. The results showed that 95.0% of conserved genes were recovered in the annotation (Supplementary Table 4).

Following gene structure annotation, we carried out functional annotation to further characterize the identified genes. Among these, 29,412 genes were successfully assigned functional information, accounting for 95.7% of the total. In particular, 8,468 genes were annotated in all five major databases, including the NCBI non-redundant protein database (Nr), UniProt, InterPro, Gene Ontology (GO), and the Kyoto Encyclopedia of Genes and Genomes (KEGG), demonstrating a high degree of functional conservation (Supplementary Figure 8,Supplementary Table 5).

In addition to protein-coding genes, annotation of the T2T genome identified four classes of non-coding RNAs, including 87 miRNAs, 450 tRNAs, 4,744 rRNAs, and 225 snRNAs (Supplementary Table 6).

### Haplotype genome annotation

Haplotype genome annotation revealed that hapA, hapB, hapC, and hapD contained 25,296, 25,395, 26,110, and 26,018 protein-coding genes, respectively (Supplementary Table 7). The completeness of these annotations, assessed by BUSCO, reached 88.0%, 87.4%, 86.8%, and 87.9% for hapA–D, respectively (Supplementary Table 8), which is fewer than the genes annotated in the T2T genome. This difference is likely due to the higher completeness of the T2T assembly, whereas the haplotype assemblies still contain a number of unresolved regions, resulting in slightly lower gene counts.

Based on haplotype annotation, each haplotype of *D. officinale* was treated as an independent genomic unit and included in a phylogenetic analysis together with other *Dendrobium* species and *Apostasia shenzhenica*. The four haplotypes (hapA–hapD) formed a tightly clustered, well-supported clade together with the published diploid *D. officinale* v3.0 genome. This *D. officinale* clade was most closely related to *D. huoshanense* and was clearly separated from other diploid *Dendrobium* species. These results support the inference that *D. officinale* is an autotetraploid (Supplementary Figure 9).

### Centromeres of the T2T *D. officinale* Genome

Although the genome assembly achieved T2T-level continuity, the prediction of centromeric regions was challenged by the complexity of highly repetitive sequences. A total of 15 centromeres were identified, with lengths ranging from 92,160 bp to 2,016,000 bp (Supplementary Table 9). For chromosomes 3, 8, 9, and 13, although centromeric regions were predicted by Centromics(27), their boundaries and features appeared ambiguous, and manual inspection indicated limited certainty; thus, these regions were labeled as “unverified”. Notably, no centromeric regions were predicted for chromosomes 10, 12, 14, and 16. This absence may reflect limitations in current prediction tools when handling highly repetitive sequences.

In addition, analysis of the relationship between centromeres and whole chromosomes showed no significant correlation between centromere size and chromosome length (Supplementary Figure 10). Based on centromere position along the chromosomes, chromosome 1, 3, 6, 8 were identified as metacentric; chromosome 4, 5, 11, 17, 19 as submetacentric; chromosome 13, 18 as acrocentric; while chromosome 2, 7, 9, 15 were characterized as telocentric (Supplementary Table 10).

Analysis of centromere composition revealed TEs enrichment in the centromeres of chromosome 3, 4, 5, 6, 8, 11, 15, and 19, whereas minisatellites predominated in those of chromosome 1, 2, 7, 9, and 17; the centromere of chromosome 18 was predominantly composed of TEs and satellite sequences. In TEs-rich centromeres, LTR retrotransposons of the Gypsy lineage constituted the dominant component, reaching 48.94% of centromeric sequence on chromosome 3. In minisatellites-rich centromeres, TEs in the centromeres of chromosome 2 and chromosome 7 were dominated by DNA transposons (Supplementary Figure 11, Supplementary Table 10, 11).

A total of 80 genes were identified within centromeres across the *D. officinale* genome, with the centromeric region of chromosome 8 containing the largest number (29 genes), whereas no genes were detected in the centromeres of chromosome 1, 15, 17, and 18. Genes located within centromeric regions exhibited low expression levels in root, stem, and leave tissues. Functional annotation was performed for 67 genes based on searches against the InterPro, NR, and UniProt databases, suggesting potential involvement of a subset of these genes in retrotranscription-related processes (Supplementary Table 12).

### Evolutionary scenario of the T2T *D. officinale* genome

Comparative genomic analysis of *D. officinale* was conducted alongside 24 representative plant species using OrthoFinder. A total of 32,210 orthogroups containing 6,373,311 genes were identified across the 25 species. Among these, 3,820 orthogroups were shared by all species, including 207 single-copy orthologous groups.

Phylogenetic analysis based on single-copy orthologous genes revealed that Orchidaceae and Poaceae diverged approximately 124.45 Mya(95% CI: 112.86–132.02 Mya). Within Orchidaceae, the genus *Dendrobium* diverged from *Phalaenopsis equestris* at approximately 39.81 Mya (95% CI: 30.84–49.70 Mya). Within the genus *Dendrobium*, *D. officinale*, *D. huoshanense*, *D. hercoglossum*, and *D. nobile* form a closely related clade. Divergence time estimates within this clade indicate that *D. officinale* and *D. huoshanense* diverged at approximately 4.61 Mya (95% CI: 3.00–6.31 Mya), while *D. hercoglossum* and *D. nobile* diverged slightly later at approximately 4.53 Mya(95% CI: 2.91–6.23 Mya) (Figure 2A).

**Figure 2.**
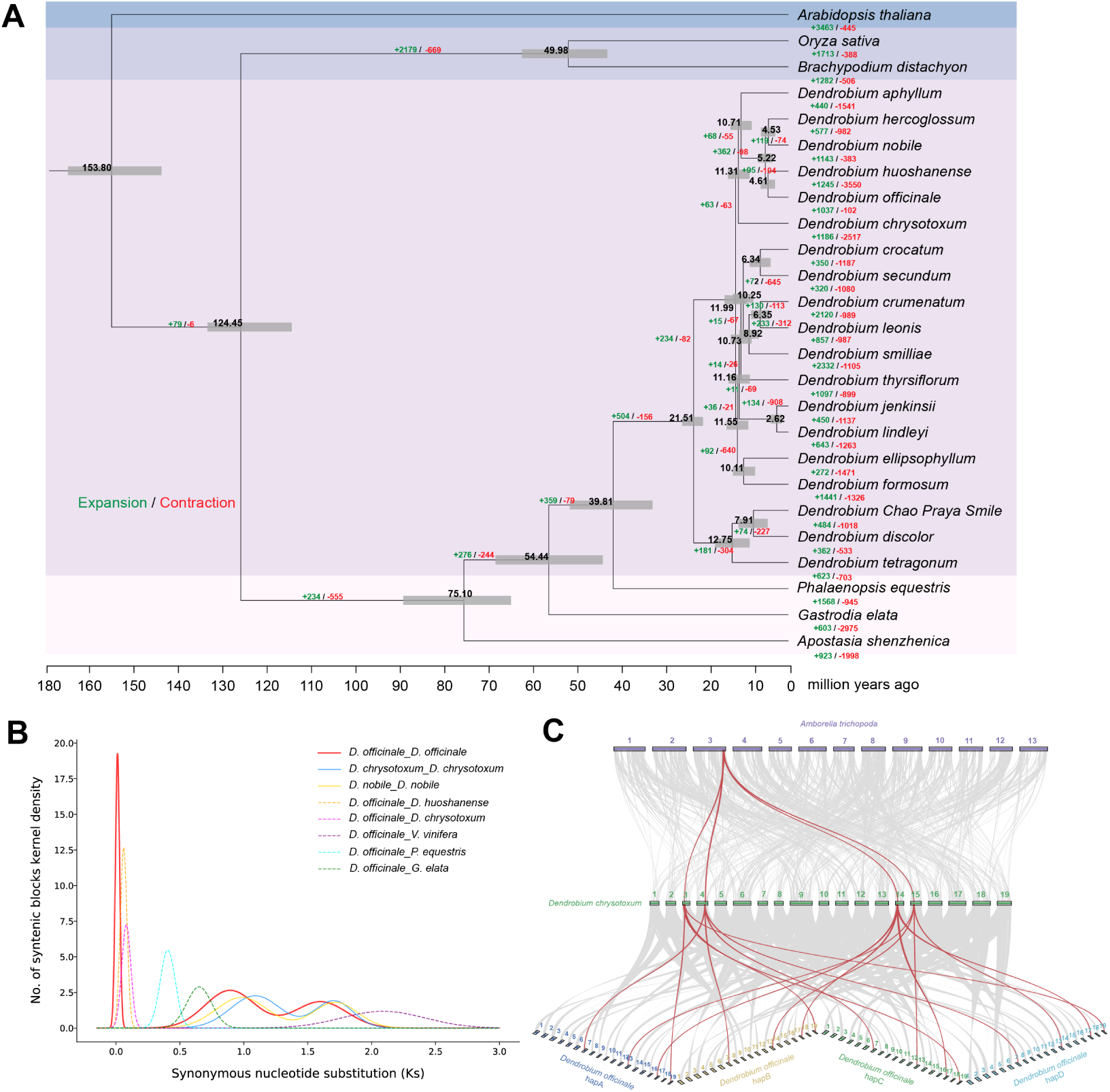
Comparative genomic analysis of *D. officinale*. (A) Phylogenetic tree reconstruction showing the evolutionary relationship between *D. officinale* and 24 representative plant species.The divergence times at internodes are indicated by black numbers, and the 95% confidence intervals of the divergence time are shown by gray bars. The numbers in green and red at branches represent the expansion and contraction of gene families, respectively. (B) Ks distribution of syntenic gene pairs. The solid red line represents the Ks distribution of paralogous syntenic gene pairs within *D. officinale*. (C) Synteny analysis of genomic regions between *Dendrobium chrysotoxum* and *Amborella trichopoda* (top), and between *Dendrobium officinale* and *Dendrobium chrysotoxum* (bottom).

In *D. officinale*, 399 species-specific orthogroups comprising 1,524 genes were detected. In the GO category, these genes were mainly enriched in peptidase-related molecular functions, including metallopeptidase, aminopeptidase and exopeptidase activities, as well as terms associated with oxidative phosphorylation and transmembrane transporter activity. KEGG pathway enrichment indicated that species-specific genes were involved in multiple metabolic and signaling pathways, such as the pentose phosphate pathway, glycolysis/gluconeogenesis, MAPK signaling pathway, and lipid-related metabolic pathways (Supplementary Figure 12).

Gene family evolution analysis showed that *D. officinale* experienced the expansion of 1,037 gene families, comprising 4,134 genes. GO enrichment analysis of genes from expanded gene families revealed significant enrichment in transporter-related molecular functions, particularly ATPase-coupled and transmembrane transporter activities, together with several methyltransferase-related terms. KEGG pathway analysis indicated that these expanded genes were mainly enriched in pathways associated with ribosome and ribosome biogenesis, photosynthesis, fatty acid metabolism, glutathione metabolism, and alkaloid biosynthesis. Taken together, these results suggest that the expansion of specific gene families in *D. officinale* may have contributed to enhanced capacities for substance transport, primary metabolism, and secondary metabolite biosynthesis, which could be related to its medicinal properties and environmental adaptability (Supplementary Figure 13).

In contrast, 102 contracted gene families, comprising 121 genes, were identified. GO enrichment analysis showed that genes from contracted gene families were primarily enriched in transmembrane transporter activity and glycosylation-related functions, including mannosyltransferase and mannan synthase activities. KEGG pathway enrichment indicated that contracted genes were mainly involved in central metabolic pathways, including ubiquinone and terpenoid–quinone biosynthesis, pentose phosphate pathway, citrate cycle, and amino acid metabolism (Supplementary Figure 14).

### Whole-genome duplication events in the tetraploid *D. officinale*

To investigate whole-genome duplication (WGD) events in the autotetraploid *D. officinale*, we calculated the synonymous substitution rates (Ks) of its genes. The Ks distribution revealed three distinct peaks at Ks = 1.60, 0.89, and 0.01, indicating that *D. officinale* has undergone at least three WGD events, including a recent WGD event (Figure 2B).

The peaks at Ks = 0.89 and Ks = 1.60 are consistent with previous findings in the genomes of Guangnan and huoshan *D. officinale*(11; 12). The Ks = 0.89 peak is shared among orchids and has been estimated to correspond to a WGD event that occurred approximately 74 Mya (90% CI: 72-78 Mya) (28). The Ks = 1.60 peak corresponds to the monocot-wide WGD event, which is estimated to have occurred around 120 Mya (95% CI: 110-135 Mya) (29). Both of these older WGD signals have also been observed in *D. nobile*, *D. chrysotoxum*, and other orchid genomes(30; 31).

Using the orchid-shared WGD event, which has been dated to approximately 74 Mya and corresponds to a Ks peak at 0.89 in *D. officinale*, we calibrated the synonymous substitution rate for *D. officinale*. Based on this calibrated rate, the most recent WGD event, characterized by a Ks peak at 0.01, was estimated to have occurred at approximately 0.86 Mya (95% CI: 0.34–1.39 Mya). This recent WGD event is further supported by comparative synteny analysis. A syntenic depth of 1:4 was observed between *A. trichopoda* and *D. chrysotoxum*, and a similar 1:4 relationship was also detected between *D. chrysotoxum* and the haplotype-resolved genome of *D. officinale*, indicating that *D. officinale* experienced an additional WGD beyond the two WGDs shared with *D. chrysotoxum* (Figure 2C, Supplementary Figure 15).

### Haplotype genome gene expression analysis

Allelic expression patterns were subsequently analyzed. A total of 12,761 tetrads were identified (Supplementary Table 13). Among them, 10,653, 9,924, and 9,859 tetrads were expressed in root, stem, and leaf tissues, respectively. Across three tissues, balanced expression patterns accounted for only 11.0%–13.3% of genes, whereas 15.9%–19.6% of genes showed suppressed expression in one haplotype (Figure 3B). Notably, genes exhibiting dominant expression in one or two haplotypes represented the largest proportion, indicating widespread allelic expression bias in the autotetraploid genome, reflecting the complexity and diversity of regulatory mechanisms underlying four-allele gene expression.

**Figure 3.**
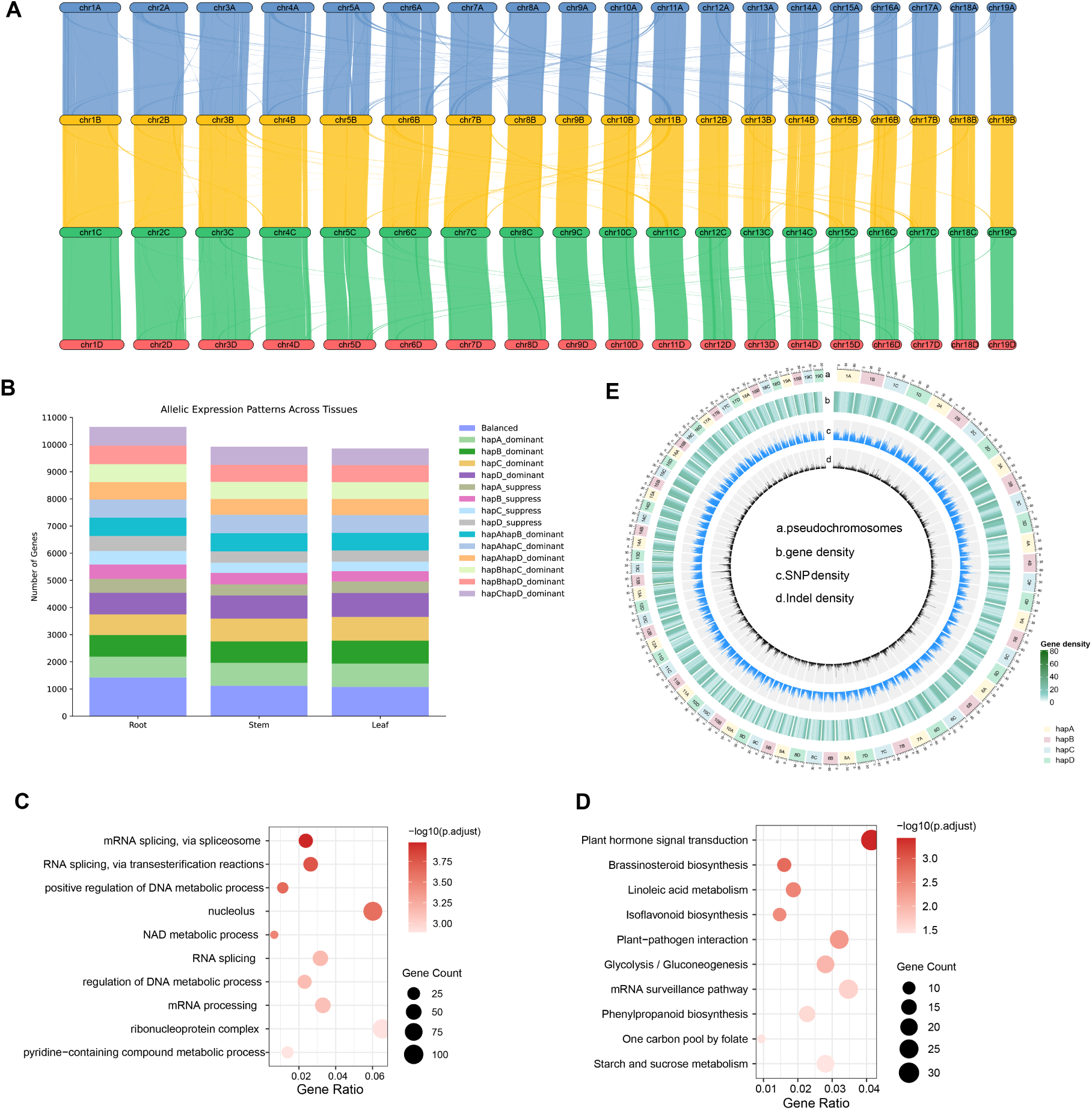
Haplotype comparison and gene expression in *D. officinale*. (A) Collinearity analysis between homologous chromosomes of *D. officinale*. (B) Expression patterns of tetra-allelic genes across root, stem, and leaf tissues. The stacked bar charts show the number of genes corresponding to each of the 15 defined expression patterns. (C) GO enrichment analysis of genes showing single-allele dominant expression in root tissue. (D) KEGG pathway enrichment analysis of genes showing single-allele dominant expression in root tissue. (E) Circos visualization of genome features in the four haplotypes of *D. officinale*. Chromosome-scale pseudomolecules (a), gene density (b), SNP density (c), and Indel density (d) are shown. For each haplotype, SNP and Indel densities were calculated using one of the other haplotypes as the reference in a circular manner: hapA vs. hapB, hapB vs. hapC, hapC vs. hapD, and hapD vs. hapA. All feature densities were computed in non-overlapping 500-kbp windows.

To explore the functional implications of tetra-allelic genes with distinct allelic expression patterns, we performed GO and KEGG enrichment analyses for four categories: dual-allele dominant, single-allele suppression, balanced, and single-allele dominant gene. The results revealed striking functional differentiation among these groups. Single-allele dominant genes were prominently enriched in RNA splicing, mRNA processing, and regulation of DNA metabolic processes, with cellular component annotations enriched in the nucleolus and ribonucleoprotein complexes, indicating a strong association with transcriptional and translational machinery. KEGG analysis further revealed significant enrichment in plant hormone signal transduction, as well as involvement in brassinosteroid biosynthesis, lipid metabolism, plant-pathogen interaction, and carbohydrate metabolism (Figure 3C,3D). Dual-allele dominant genes were enriched in RNA and tRNA methylation, protein glycosylation, and vesicle-mediated transport, indicating roles in modulating post-transcriptional modifications and intracellular trafficking. Single-allele suppression genes were primarily associated with hormone signaling, defense responses, and lipid metabolism, reflecting their potential role in the fine-tuning of stress responses and secondary metabolism. Balanced genes in roots showed enrichment in vesicle-related structures, implying functions in cellular homeostasis and trafficking (supplementary Figure 16). Overall, these findings reveal that allele expression patterns in *D. officinale* are functionally partitioned, potentially contributing to regulatory complexity in this autotetraploid genome. regulation may contribute to phenotypic plasticity and environmental adaptation in *D. officinale*.

### Haplotype genomic variation analysis

Synteny analysis indicated that the four homologous chromosome groups maintain a high degree of collinearity, preserving a strict 1:1 syntenic correspondence (Figure 3A).

To further investigate the genetic variation among these haplotypes, pairwise comparisons were conducted to systematically characterize the distribution of single nucleotide polymorphisms (SNPs), insertions and deletions (Indels), and structural variations (SVs) (Figure 3E, Supplementary Table 14). SNP counts were relatively consistent across haplotype pairs, ranging from 6,300,439 to 6,357,982, suggesting similar levels of single-nucleotide variation among the four haplotypes. In contrast, Indel counts were slightly more variable, ranging from 1,303,678 to 1,337,717. Despite these differences in count, the total length of Indels remained relatively stable across all haplotype pairs, varying only slightly from 3.32 to 3.40 Mbp. Structural variations showed similar patterns, with counts ranging from 24,525 to 25,186. The total size of SVs varied between 45.90 Mbp and 47.48 Mbp, with the hapC–hapD pair exhibiting the greatest extent of structural variation. Comparisons with the *D. officinale* v3.0 genome revealed slightly higher variation, with SNPs ranging from 7.74 to 7.82 million and SV sizes of 54.8–56.3 Mbp.

Collectively, these results indicate that despite widespread sequence variation, the four *D. officinale* haplotypes maintain an overall conserved genome structure. This combination of high collinearity and controlled variation suggests evolutionary stability of chromosome architecture alongside functional diversification at the sequence level.

### Differential gene expression across tissues

Transcriptomic profiling of roots, stems, and leaves of *D. officinale* revealed distinct expression patterns among different tissues. After quality control, all nine RNA-seq datasets exhibited high sequencing quality, with Q20 values ranging from 97.53% to 98.09% (Supplementary Table 1). Principal component analysis (PCA) and hierarchical clustering showed that samples clustered primarily by tissue type, with higher correlations within tissues than between tissues, indicating strong biological reproducibility (Supplementary Figure 17).

Differential expression analysis identified 3,258, 2,980, and 4,871 differentially expressed genes (DEGs) in the stem vs root, stem vs leaf, and leaf vs root comparisons, respectively (| log_2_ FC| > 1, Padj < 0.05). Among these, 634 DEGs were shared by all three pairwise comparisons (Supplementary Figure 18). To identify tissue-specific expression patterns, three sets of tissue-upregulated genes were extracted based on the pairwise DEGs, as described in the Methods. This analysis yielded 1,303 root-upregulated genes, 331 stem-upregulated genes, and 974 leaf-upregulated genes.

Leaf-upregulated genes were predominantly enriched in GO terms associated with chloroplast organization, thylakoid membranes, and photosynthesis-related biological processes. Consistently, KEGG pathway analysis revealed significant enrichment in photosynthesis, carbon fixation, porphyrin metabolism, and circadian rhythm–related pathways. This indicates that the leaf-upregulated genes are enriched in photosynthesis, light signal reception and conversion, as well as related basic metabolic processes.

Stem-upregulated genes were mainly associated with developmental processes, including meristem development, regulation of growth, and responses to light stimuli at the GO level. KEGG enrichment further indicated the involvement of carbohydrate metabolism, amino acid metabolism, and terpenoid biosynthesis pathways. These results suggest that stem-upregulated genes are functionally associated with developmental regulation and specific metabolic processes, including carbohydrate and terpenoid metabolism.

Root-upregulated genes showed significant enrichment in GO terms related to secondary metabolic processes, hormone regulation, and glycosyltransferase activities. Correspondingly, KEGG analysis highlighted pathways involved in flavonoid and phenylpropanoid biosynthesis, plant hormone signal transduction, and ABC transporters (Supplementary Figure 19). These results suggest that root-upregulated genes are involved in secondary metabolite biosynthesis, hormone signaling, and transport, which may contribute to the root’s roles in environmental adaptation, defense, and metabolite distribution.

### Variation of *SWEET* gene repertoires among *Dendrobium* species

The SWEET family represents a class of sugar transporters widely distributed in plants. *SWEET* genes play a central role in facilitating the transmembrane transport of mono- and disaccharides, especially sucrose, glucose, and fructose (32). SWEET proteins exhibit substrate selectivity and are involved not only in sugar transport but also in plant growth, development, and responses to biotic and abiotic stresses(33; 34).

We identified the *SWEET* genes in the genomes of *D. officinale* and 18 other *Dendrobium* species, and constructed a phylogenetic tree based on these data (Figure 4A, Supplementary Table 15). These species belong to 12 different sections. Among them, six species are tree epiphytic, two are rock epiphytic, and ten have mixed epiphytic habits, growing on both trees and rocks. Most species are distributed in tropical regions, while *D. huoshanense* and *D. officinale* are found in subtropical areas, and *D. nobile* spans both tropical and subtropical zones (Supplementary Table 16).

**Figure 4.**
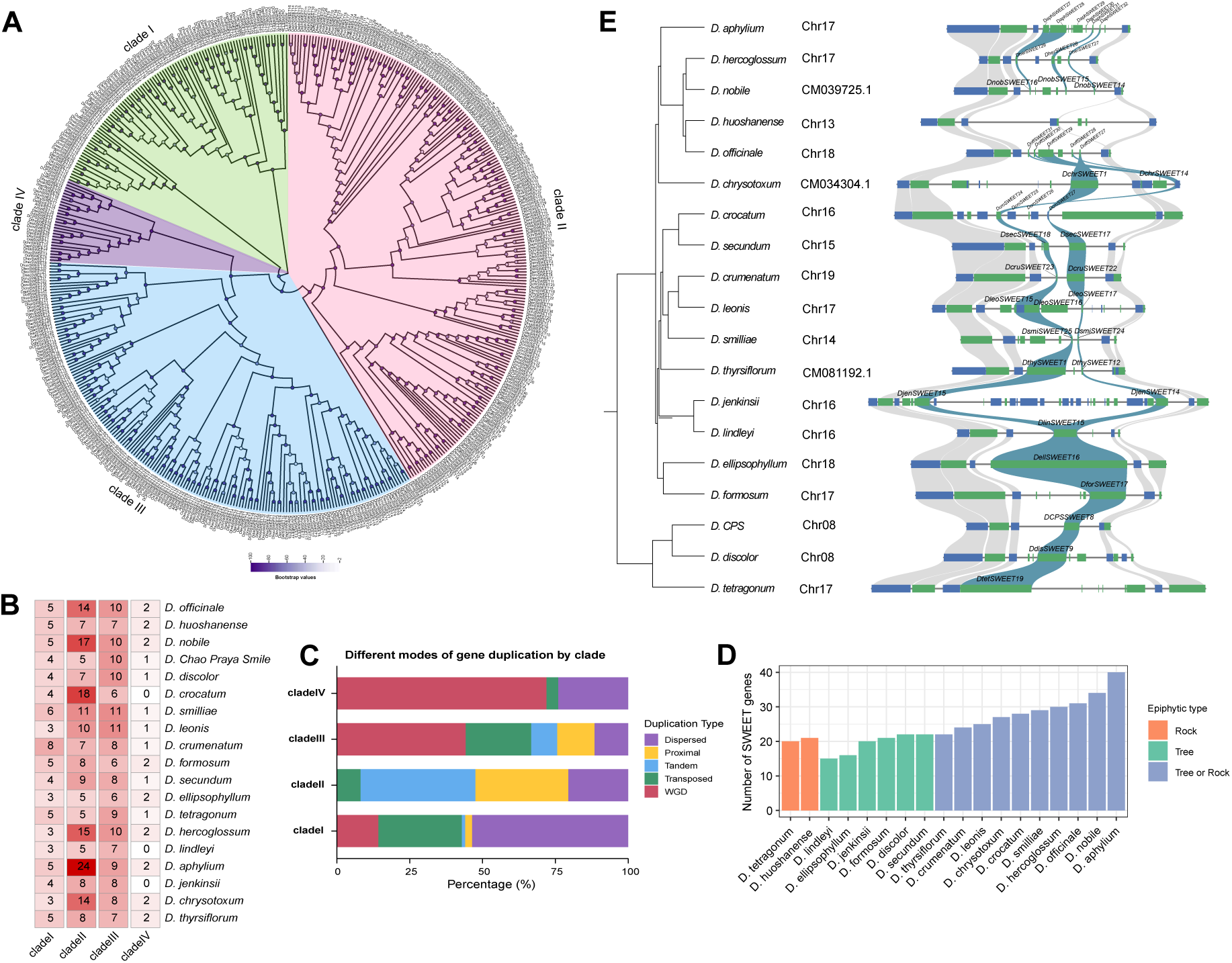
Comparative analysis of the SWEET gene family in the *Dendrobium* genus. (A) A phylogenetic tree of SWEET proteins from 19 *Dendrobium* species, with those from *Arabidopsis thaliana* and *Oryza sativa*, showing their classification into four clades (clade I, green; clade II, pink;clade III, blue; clade IV, purple). (B) The distribution of *SWEET* gene numbers across the studied *Dendrobium* species. (C) Statistics of gene duplication modes for the *SWEET* genes within the four clades. (D) Comparison of *SWEET* gene numbers among *Dendrobium* species with different epiphytic types. The hybrid *D. Chao Praya Smile* was excluded from comparative analyses because its epiphytic type and ecological background remain unclear. (E) Microsynteny analysis of *SWEET* genes across *Dendrobium* species. Gene names are indicated, and syntenic *SWEET* genes are highlighted in cyan.

The number of *SWEET* genes in these species was observed to range from 15 to 40, with *D. lindleyi* having the fewest, possessing only 15 *SWEET* genes, while *D. aphylium* had the highest number, with a total of 40 *SWEET* genes. The greatest variation in gene number was observed in clade II, where four species (*D. ellipsophyllum*, *D. CPS*, *D. tetragonum*, and *D. lindleyi*) each contain only 5 genes, while *D. aphylium* contains 24 genes. Members of clade IV are relatively few, usually no more than 2 genes, and are absent in *D. crocatum*, *D. jenkinsii*, and *D. lindleyi*, suggesting potential gene loss during evolution (Figure 4B). Furthermore, we analyzed the duplication patterns of *SWEET* genes from *Dendrobium* species within the four clades of the phylogenetic tree. The expansion of clade III and clade IV was primarily driven by whole-genome duplication, whereas clade II showed substantial contributions from tandem, proximal, and dispersed duplications. In contrast, clade I was mainly expanded through transposed and dispersed duplications (Figure 4C).

Interestingly, species with mixed epiphytic habits generally possess a higher number of *SWEET* genes, while those with a single epiphytic habit tend to have fewer. This suggests that the epiphytic habit may influence the number of *SWEET* genes, possibly due to differences in ecological adaptation strategies (Figure 4D). Additionally, a correlation was observed between *SWEET* gene number and plant size. For example, in section *Dendrobium*, *D. huoshanense*—a smaller plant—has fewer *SWEET* genes, while the larger species, such as *D. officinale*, *D. nobile*, and *D. aphylium*, possess 31, 34, and 40 *SWEET* genes, respectively. A similar trend was observed in section *Densiflora* (Supplementary Table 16).

In *Dendrobium* species, we identified a syntenic *SWEET* gene tandem (Figure 4E), within which a subset of genes exhibit clear collinear relationships across species, suggesting the presence of an ancestral *SWEET* locus. Extensive lineage-specific tandem duplications within this block have led to substantial variation in *SWEET* gene copy number and structure among species. Gene duplication analysis showed that copy numbers within this block vary from 1 to 6 among *Dendrobium* species. Notably, no *SWEET* gene copies were detected in this block in *D. huoshanense*, which may reflect lineage-specific gene loss or limitations in genome assembly and annotation. In addition to copy number variation, *SWEET* genes within this syntenic block exhibit substantial length variation. Several genes display unusually expanded gene structures, which are largely associated with the presence of ultra-long introns (Supplementary Figure 20). Such structural expansions could potentially be related to transposable element insertions or other genomic rearrangement events, which may contribute to increased structural and functional diversity. However, the precise evolutionary mechanisms underlying these structural variations remain to be further investigated.

In *Dendrobium*, SWEET gene family size is associated with ecological adaptation, as species with mixed epiphytic habits or larger plant size generally possess more *SWEET* genes than those restricted to a single epiphytic niche or smaller growth forms. Building on this genus-level perspective, we next focused on *D. officinale* to investigate the composition, distribution, and expression patterns of its SWEET gene family.

### SWEET genes in D. officinale

In this study, using newly assembled T2T genome we performed a genome-wide identification of the SWEET gene family in *D. officinale*. We identified 31 *DoffSWEET* genes and assigned standardized names based on their chromosomal positions. The 31 *DoffSWEET* genes are distributed across ten chromosomes; chromosome 11 contains six *DoffSWEET* genes, representing the highest density of *SWEET* genes in the assembly. Subcellular localization prediction indicated that 28 *DoffSWEET* proteins are localized to the cell membrane and 3 are localized to the chloroplast (Supplementary Table 17).

Phylogenetic analysis placed the 31 genes into four clades: 5 genes in clade I, 14 in clade II, 10 in clade III, and 2 in clade IV. Previous studies have reported that clade I and II SWEET proteins are generally associated with hexose transport, clade III members primarily mediate sucrose transport, and clade IV members are typically involved in fructose transport (32). Furthermore, clade II and III in *D. officinale* also exhibited expansion driven by multiple duplication modes, whereas clade IV was relatively conserved, containing only two WGD-derived genes (Figure 5A).

**Figure 5.**
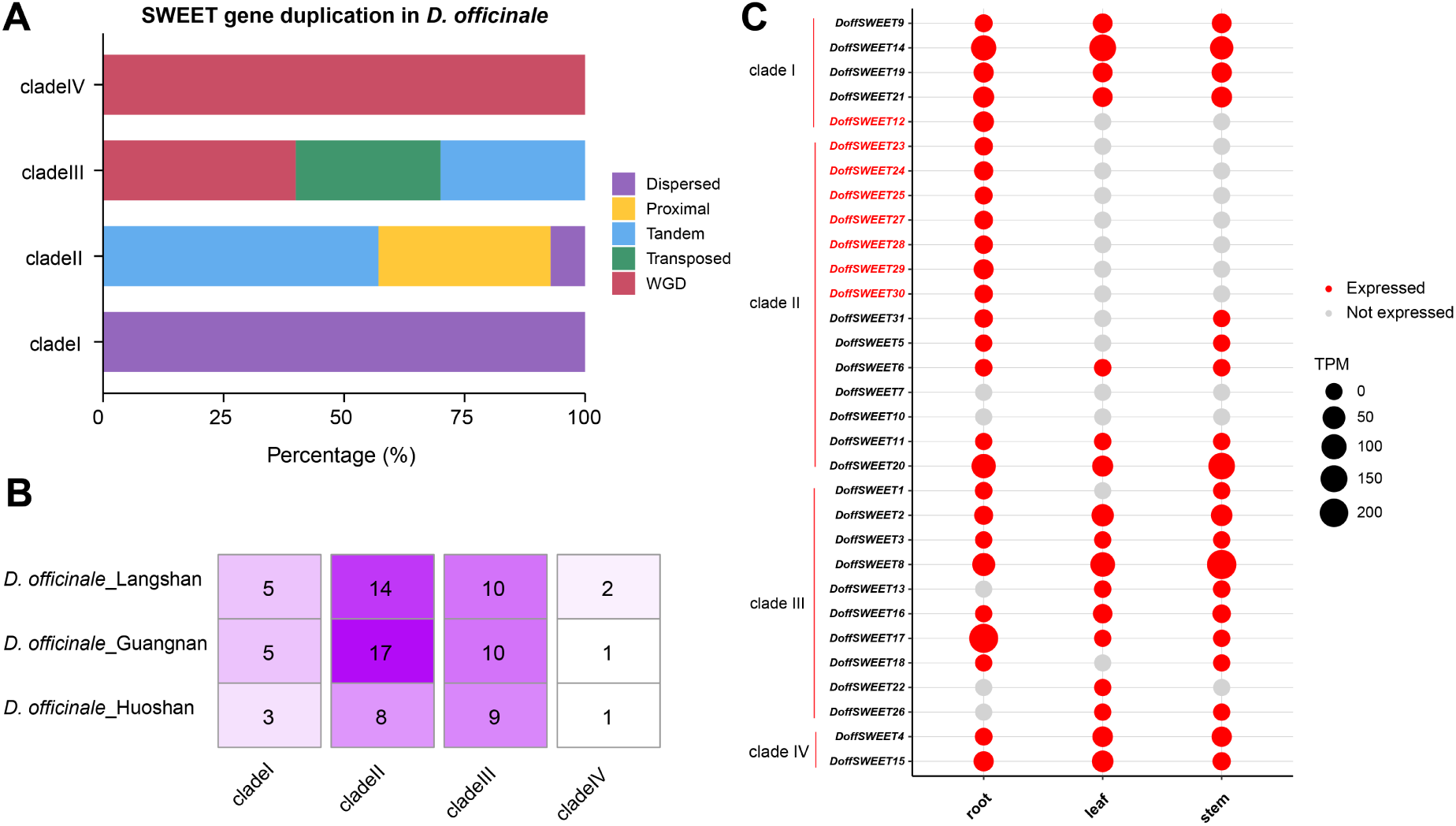
Characterization of the SWEET gene family in *D. officinale*. (A) Duplication modes of *SWEET* genes in the Langshan *D. officinale*(this study). (B) Comparison of *SWEET* gene numbers among the Guangnan, Huoshan, and Langshan *D. officinale*. The Huoshan *D. officinale* contains the fewest *SWEET* genes, with the most pronounced difference observed in clade II. (C) Expression profiles of *SWEET* genes in root, stem, and leaf tissues of the Langshan *D. officinale*. Eight *DoffSWEET* genes, highlighted in red, were specifically expressed in roots.

Comparison of *D. officinale* accessions from Guangnan, Langshan, and Huoshan revealed differences in the number of *SWEET* genes. The Guangnan accession contains 33 *SWEET* genes, the Langshan accession contains 31, and the Huoshan accession contains 21. At the clade level, the Guangnan accession exhibits 5, 17, 10, and 1 members in clades I–IV, respectively; the Langshan accession exhibits 5, 14, 10, and 2 members; and the Huoshan accession exhibits 3, 8, 9, and 1 members. The most pronounced difference occurs in clade II, which is less numerous in the Huoshan accession than in Guangnan and Langshan(Figure 5B). These results indicate that the expansion and contraction of the SWEET gene family occur not only among species but also within a single species across different geographic accessions. Even within the same species, the number of *SWEET* genes can vary substantially, reflecting the influence of local environments and ecological conditions.

In the Langshan accession analyzed in this study, eight *DoffSWEET* genes were expressed exclusively in roots, with no detectable expression in stems or leaves (Figure 5C). Among them, seven belonged to clade II and one to clade I. Notably, within clade II, *DoffSWEET23*, *DoffSWEET24*, and *DoffSWEET25* exhibited a proximal duplication pattern, whereas *DoffSWEET27*, *DoffSWEET28*, *DoffSWEET29*, and *DoffSWEET30* were tandemly duplicated and located within the syntenic block shown in Figure 4E. This distribution suggests that the proximal duplicates may have retained ancestral functions, while the tandemly duplicated genes likely arose from recent local expansions within a conserved genomic region, potentially allowing functional diversification. Considering that roots are the primary site of orchid mycorrhizal symbiosis in *D. officinale*, the strict root-specific expression of these genes, together with their duplication patterns and conserved genomic context, suggests that they may have undergone functional specialization associated with root biology and may contribute to the molecular processes underlying root mycorrhizal symbiosis.

### Cis-acting element of *DoffSWEET* gene

We analyzed the types and numbers of cis-acting elements in the promoter sequences of *D. officinale SWEET* genes (Supplementary Figure 21). Among these, abiotic and biotic stress-responsive elements were the most abundant, with a total of 468 identified in the 31 *SWEET* genes; light-responsive elements accounted for 309, phytohormone-responsive elements for 216, and elements related to plant growth and development were the least abundant, with 42.

This pattern was also observed in the eight root-specific *DoffSWEET* genes. Specifically, *DoffSWEET14*, *DoffSWEET23*, *DoffSWEET27*, *DoffSWEET28*, *DoffSWEET29*, and *DoffSWEET30* contained the highest number of abiotic and biotic stress-responsive elements in their promoters, whereas *DoffSWEET24* and *DoffSWEET25* had the most light-responsive elements, followed by abiotic and biotic stress-responsive elements. Overall, these results suggest that the promoters of these root-specific genes possess regulatory potential to respond to environmental stresses and biotic signals.

## Discussion

In this study, a reference-quality telomere-to-telomere genome assembly of *Dendrobium officinale* is reported for the first time, substantially improving on earlier versions and enabling deeper genomic insights. For this T2T genome, all 38 telomeres were successfully identified, and 15 centromeric regions were fully characterized. This comprehensive, gap-free genome assembly facilitates highly accurate structural and functional genome annotation, providing a solid foundation for in-depth comparative genomics, evolutionary analysis, and functional genomics studies. To our knowledge, this is also the first T2T genome assembly in the Orchidaceae family, representing a significant milestone in orchid genomics with the first gap-free chromosomal reference for the family.

Notably, four haplotype-resolved assemblies were generated, capturing the allelic diversity inherent to the tetraploid nature of *D. officinale*. This work also provides the first genome resource for a tetraploid *Dendrobium* species, whereas previously published genomes in the genus were limited to diploid representatives. We identified 12,761 four-allele loci and explored their diverse expression patterns. Comprehensive analyses of genomic variations among haplotypes were also conducted. These results provide an invaluable genomic resource for investigating allele-specific expression, haplotype interactions, and gene dosage effects—key factors in understanding trait regulation and adaptation in polyploid species.

We inferred that the most recent whole-genome duplication in tetraploid *D. officinale* corresponds to a relatively recent autotetraploidization event that occurred approximately 0.86 Mya. However, it should be noted that dating autopolyploidization events based on Ks distributions implicitly assumes little to no sequence divergence among haplotypes at the time of genome duplication. In natural populations, the diploid progenitors of autotetraploid species often retain a certain level of heterozygosity; consequently, the homoeologous gene copies generated by the duplication event are not entirely identical. Under these circumstances, Ks-based time estimates may partially reflect divergence that predated the polyploidization event, and therefore are more likely to represent an upper bound for the timing of the autotetraploidization.

Within the genus *Dendrobium*, the number of *SWEET* genes varies significantly, ranging from approximately 15 to 40, and is closely associated with epiphytic growth habit. This suggests that the size of the SWEET gene family may be linked to the environmental adaptability of these plants. Similar patterns have been reported in other species. For instance, in *Brassica rapa*, *SWEET* genes have been extensively retained and expanded following whole-genome triplication, and these expanded genes exhibit differential expression across tissues and under stress conditions, indicating potential functional diversification in response to abiotic stresses (35). In hexaploid wheat, *SWEET* genes are widely activated under drought and salt stress, further supporting the notion that *SWEET* gene expansion may contribute to plant adaptation to complex environments (36). Taken together, these observations suggest that variation in SWEET gene number within *Dendrobium* may play an important role in ecological adaptation.

Variation in SWEET gene number is also observed among *D. officinale* from different geographic locations: Guangnan *D. officinale* harbors 33 *SWEET* genes, Langshan *D. officinale* has 31, and Huoshan *D. officinale* contains 21. *D. officinale* from Langshan grows on the cliffs of Danxia landforms, where the surface temperature is high and humidity is low during summer, imposing substantial environmental stress. The increased number of *SWEET* genes in Langshan *D. officinale*, particularly the expansion of clade II members, may be associated with maintaining efficient sugar allocation under adverse conditions. In contrast, Huoshan *D. officinale* has fewer *SWEET* genes, possibly reflecting the lower environmental stress and reduced demand for internal sugar transport. *SWEET* genes fine-tune sugar distribution across different tissues and cellular compartments, potentially functioning as osmoprotectants and supporting cellular activity, thereby contributing to plant survival under drought, high salinity, low temperature, or high temperature stress (36; 37; 38; 39; 40; 41).

The growth and development of *D. officinale* depend on its association with orchid mycorrhizal fungi (OMF). Previous studies have shown that *SWEET* genes, which encode sugar transporters, are involved in sugar transport and allocation during mycorrhizal interactions in various plant species. For example, *GmSWEET6* in soybean participates in sucrose transport during arbuscular mycorrhizal interactions, whereas *MtSWEET1b* in Medicago truncatula is strongly upregulated in arbuscule-containing cells and primarily functions as a glucose transporter, facilitating nutrient flow at the plant–fungus interface (42; 43). In Guangnan *D. officinale*, transcriptome analysis revealed that a *SWEET* gene was significantly upregulated in roots colonized by OMF, suggesting its potential role in regulating sugar allocation and contributing to the establishment and maintenance of plant–fungus associations (44). In this study, we identified eight *DoffSWEET* genes that were specifically expressed in roots, which are the primary sites of orchid mycorrhizal colonization and symbiosis. Given that orchids generally establish asymmetric, deceptive associations with fungi, these root-specific *SWEET* genes are likely to regulate sugar partitioning within root tissues, thereby enhancing the ability of *D. officinale* to attract and maintain fungal colonization(45; 46). Such sugar-regulatory mechanisms may represent the molecular basis of the deceptive symbiotic strategy of *D. officinale* and its ecological adaptation to nutrient-poor epiphytic environments.

Future research could investigate the expression patterns of *SWEET* genes in *D. officinale* across different geographical regions and under varying environmental conditions, thereby providing further insights into their roles in sugar allocation and responses to abiotic stresses. Moreover, considering the functional studies involving gene knockout or overexpression are still not possible at present, eight *DoffSWEET* genes that were specifically expressed in roots of tetraploid *D. officinale* in Langshan area can be treated as an especial system to elucidate the roles of root-specific *SWEET* genes in mediating sugar transport between plants and mycorrhizal fungi when comparing with other populations in which the roots have different number of *DoffSWEET* genes. Such studies will contribute to a deeper understanding of the relationship between the SWEET gene family and ecological adaptation, offering theoretical foundations for the conservation and cultivation of *D. officinale*

In summary, these results provide a comprehensive view of the genomic architecture, genome evolution, and biosynthetic potential of *D. officinale* (Langshan, Hunan). The integration of telomere-to-telomere assembly and haplotype-resolved analysis enhances genome continuity and allelic resolution, enabling accurate annotation and comparative analyses in this complex tetraploid *Dendrobium* species. This high-quality genomic resource establishes a critical foundation for functional genomics and evolutionary research in *Dendrobium*. Moreover, our identification and characterization of the SWEET gene family, particularly the root-specific candidates, point to their potential roles in mediating sugar transport during orchid mycorrhizal symbioses.

## Methods

### Sampling and whole-genome sequencing

Fresh samples of *Dendrobium officinale* (cultivar ID: LS01) were collected from its cultivation site on Langshan Mountain, Shaoyang, Hunan Province, China. Genomic DNA was extracted from fresh leaves using the CTAB method. The quality and concentration of the extracted DNA were assessed using 0.75% agarose gel electrophoresis, a NanoDrop One spectrophotometer (Thermo Fisher Scientific), and a Qubit 3.0 Fluorometer (Life Technologies, Carlsbad, CA, USA).

Paired-end sequencing was performed on the DNBSEQ-T7 platform. Raw reads were filtered using fastp v0.23.4(47), discarding those with more than 40% of bases below a quality score of Q20. The resulting clean reads were used for downstream analyses.

Hi-C libraries were prepared from fresh leaves using in situ crosslinking with formaldehyde, followed by a standard Hi-C protocol to capture chromatin conformation. The libraries were sequenced on the DNBSEQ-T7 platform using 150 bp paired-end reads.

For Oxford Nanopore sequencing, libraries were prepared using the SQK-LSK110 ligation sequencing kit according to the manufacturer’s standard protocol. The purified libraries were loaded onto R9.4 Spot-On Flow Cells and sequenced on the PromethION platform (Oxford Nanopore Technologies, Oxford, UK) for 48 hours at Wuhan Benagen Technology Co., Ltd. (Wuhan, China). Base calling was conducted using Guppy (Oxford Nanopore Technologies, https://nanoporetech.com/).

For PacBio HiFi sequencing, high-molecular-weight genomic DNA was extracted from fresh leaves and used to construct circular consensus sequencing (CCS) libraries with a targeted insert size of approximately 15 kbp. Libraries were sequenced on the PacBio Sequel II platform to generate HiFi reads for de novo genome assembly.

### Transcriptome sequencing

Total RNA was extracted from roots, stems, and leaves of *D. officinale*, with three biological replicates per tissue. Sequencing was performed on the DNBSEQ-T7 platform (MGI) using the cPAS sequencing strategy at BGI.

Additionally, total RNA was extracted from the leaves of *Dendrobium officinale* using the RNAprep Pure Plant Plus Kit according to the manufacturer’s instructions (Tiangen Biotech, Beijing, China). The RNA samples were pooled and subjected to cDNA library construction using a strand-switching method with the cDNA-PCR Sequencing Kit (SQK-PCS109). Sequencing was performed on the PromethION platform (Oxford Nanopore Technologies, Oxford, UK).

### Genome size estimation

Flowers of *D. officinale* were collected and subsequently sent to Jindi Future Biotechnology Co., Ltd. (Beijing, China) for flow cytometry analysis. Diploid Solanum lycopersicum (tomato) was used as an internal reference standard to estimate the nuclear DNA content.

Additionally, k-mer analysis was performed using short-read data. K-mers were counted with Jellyfish v2.3.1(48), and the k-mer frequency distribution was analyzed using GenomeScope v2.0(49). The analysis showed a higher abundance of the aaab k-mer pattern compared to aabb, a pattern often observed in autotetraploid genomes. Based on the k-mer profile, the genome size of autotetraploid *D. officinale* was estimated to be approximately 1.1 Gbp.

### Genome assembly and pseudochromosome construction

A de novo genome assembly was generated using hifiasm v0.21.0 with the parameters --n-hap 4 --hg-size 1.1g (19), integrating PacBio HiFi reads, ultra-long Oxford Nanopore Technologies (ONT) reads and Hi-C data. This initial assembly produced 1,635 contigs. Redundant sequences were removed using Purge dups v1.2.5 (50), yielding 130 non-redundant contigs.

Contig-level polishing was performed using NextPolish v1.4.1 (51). Three rounds of polishing were conducted using ONT reads (including ultra-long reads), followed by three additional rounds using PacBio HiFi reads. Subsequently, four rounds of polishing were carried out using DNBSEQ-T7 short reads to further correct residual base-level errors.

Hi-C reads were aligned to the polished contigs using BWA-MEM v0.7.18 (20), and a preliminary chromosome-scale assembly was constructed with HapHiC v1.0.5 (21), achieving an anchoring rate of 99.99%. Scaffold structure, chromosomal boundaries and potential misassemblies were manually inspected and corrected using Juicebox (22), resulting in 19 pseudochromosomes. Remaining assembly gaps (*n* = 140) were closed using TGS-GapCloser v1.2.1 (23) with both ONT and HiFi reads.

To complete telomeric regions, all ONT reads were aligned to the assembly using Winnowmap v2.0 (52). Reads mapping within 100 bp of chromosome termini were extracted and locally reassembled into consensus telomeric sequences using Medaka v2.0.1. These consensus sequences were aligned back to the assembly using minimap2 v2.24 (53), and the best-supported alignments were used to replace terminal regions. A final round of polishing was performed using NextPolish2 v0.2.0 (54), resulting in a gap-free, telomere-to-telomere collapsed assembly.

### Haplotype-resolved assembly

Obtaining chromosome-scale phased assemblies for a tetraploid genome is inherently challenging because highly similar homologous chromosomes and abundant repeats can cause contig fragmentation, haplotype switching and cross-homolog mis-joins when phasing and scaffolding are performed in a single step. We initially explored a direct hifiasm-based haplotype-assembly workflow to generate four phased assemblies followed by standard post-processing (purging redundancy, scaffolding, gap closing and polishing). However, the resulting assemblies were highly fragmented and showed unstable haplotype completeness, making them unsuitable for downstream haplotype-specific analyses. We therefore adopted a conservative two-stage strategy that prioritizes haplotype enrichment at the read/contig level and chromosome-scale structural consistency at the scaffolding stage.

First, PacBio HiFi reads were partitioned into four haplotype-enriched read sets based on SNP information. HiFi reads were aligned to the collapsed assembly using minimap2 v2.24, variants were called using FreeBayes v1.3.8 (55), and polyploid phasing was performed using WhatsHap Polyphase v2.3 (56) to assign reads into four haplotype-enriched subsets.

Each haplotype-enriched read set was independently assembled using hifiasm, producing four haplotype-specific contig assemblies. Redundant sequences were removed using Purge dups v1.2.5, and contigs were polished with three rounds of NextPolish v1.4.1 using HiFi reads.

To obtain chromosome-scale haplotype assemblies, Hi-C data were used to infer long-range contig order and orientation and to correct potential structural mis-joins, while the primary haplotype separation was achieved at the read/contig level as described above. Hi-C reads were mapped to each haplotype contig set and used for scaffolding to generate chromosome-scale scaffolds. Scaffold structures were manually inspected and adjusted in Juicebox, with reference to the chromosomal organization of the collapsed pseudochromosomes, to ensure that each haplotype assembly follows a consistent 19-chromosome framework. The same gap-closing strategy described above was applied to each haplotype assembly.

Collectively, this procedure yields haplotype-resolved pseudochromosomes that retain haplotype-specific sequence variation while maintaining a chromosome-scale structure consistent with the collapsed assembly.

### Repeat annotation

Repetitive sequences were annotated by combining homology-based and de novo prediction approaches. First, long terminal repeat (LTR) retrotransposons in the *D. officinale* genome were identified using two tools: LTR FINDER v1.0.7(57) and LTRharvest v1.6.5(58). The LTR candidates predicted by these two tools were subsequently integrated and filtered using LTR retriever v2.9.8(59) to remove redundancy and produce a high-confidence, non-redundant set of intact LTR sequences.

Subsequently, RepeatModeler v2.0.3(60) was employed to perform de novo prediction of repetitive elements based on the genome sequence. The results from both LTR retriever and RepeatModeler were merged to construct a de novo repeat library. This library was then combined with known repeat sequences from related species in the Repbase database to form a comprehensive repeat library, which was used by RepeatMasker v4.1.2(61) to annotate repetitive elements across the genome.

The identification and classification of satellite repeats were performed following the pipeline described in the telomere-to-telomere assembly of the maize genome(62). Satellite repeats were identified using Tandem Repeats Finder v4.09(26). Tandem repeats with copy numbers less than five were filtered out. The remaining repeats were categorized based on the length of their repeat units: repeats with unit lengths less than 10 bp were defined as microsatellites, those between 10 and 100 bp as minisatellites, and those longer than 100 bp as satellites.

### Genome annoation

Gene structure annotation of T2T *D. officinale* was performed by integrating three strategies: transcriptome-based prediction, homology-based prediction, and de novo prediction.

For transcriptome-based prediction, RNA-Seq data were obtained from nine samples, including three biological replicates each from root, stem, and leaf tissues. The raw reads were filtered using fastp, which includes automatic adapter removal and discards reads containing more than 40% of bases with quality scores below Q15. The cleaned reads were then aligned to the *D. officianle* genome using HISAT2 v2.0.3(63). Full-length ONT transcriptome reads were processed using NanoFilt v2.8.0(64)with the following parameters: -q 10 --headcrop 30 --minGC 0.3 to remove low-quality sequences, and full-length transcripts were identified using Pychopper v2.7.2(65). The identified full-length reads were subsequently aligned to the reference genome using minimap2 v2.24. StringTie v2.2.3 (66) was then employed to reconstruct transcripts by integrating the alignment results from both short-read and long-read data(parameter: –mix). Finally, TransDecoder v5.7.1(67) was used to predict coding regions from the reconstructed transcripts, resulting in the identification of candidate coding genes.

For homology-based prediction, protein sequences from five species — *D. nobile*, *D. chrysotoxum*, *D. thyrsiflorum*, *D. catenatum*, and *P. equestris* — were aligned to the T2T *D. officinale* genome using tblastn. Protein sequences with less than 90% identity were filtered out. The remaining aligned sequences were used for transcript and coding region prediction with miniprot v0.12(68).

For de novo prediction, Augustus v3.5(69), Helixer v0.3.4(70), and GeneMark v4.7.1(71) were used to annotate genes based on the repeat-masked *D. officinale* genome. Augustus was trained using models generated from BUSCO results, Helixer performed cross-species gene prediction using deep learning, and GeneMark employed an unsupervised algorithm that does not require a training set.

The results from transcriptome-based, homology-based, and de novo prediction were integrated using EVM v2.1.0. Subsequently, PASA v2.5.3(72) was utilized to refine the gene models by incorporating transcriptome evidence, correcting exon-intron boundaries, adding UTRs, and modeling alternative splicing events, followed by the removal of extremely short genes and those containing premature stop codons to improve annotation quality. The final gene set was evaluated using BUSCO to assess completeness.

Gene function annotation is the process of predicting the function of target proteins by aligning their sequences with known functional protein sequences in databases. The methods for gene function annotation can be divided into two main approaches: the first is based on sequence homology, where the protein sequences of *D. officinale* are aligned with the UniProt, Nr, GO, and KEGG databases using BLASTP to obtain related functional information and potential involvement in metabolic pathways; The second is based on domain similarity, where the protein sequences of *D. officinale* were analyzed using InterProScan v5.51(73) to identify conserved domains, assign protein families, and predict functional site.

### Non-coding RNAs annotation

Non-coding RNAs are RNA molecules that do not encode proteins, including rRNA, tRNA, snRNA, snoRNA, and miRNA, among others. These RNAs perform their biological functions directly after transcription, without the need for translation into proteins. To predict the locations of these non-coding RNAs, we employed different tools: barrnap v0.9(74) for rRNA prediction in the genome, tRNAscan-SE v2.0.9(75) for tRNA prediction, and Infernal v1.1.5(76)for the prediction of snRNA, miRNA, and other non-coding RNAs.

### Centromere Characterization

Centromeric regions were identified using Centromics v0.3(27), integrating genome sequence data, HiFi reads, ONT reads and Hi-C data. The relationship between centromere length and chromosome length was evaluated. Chromosomes were classified as metacentric, submetacentric, acrocentric, or telocentric based on relative centromere position and arm ratio (long-arm length/short-arm length)(77). To characterize the genomic landscape of centromeres, transposable elements and tandem repeat sequences were extracted from centromeric intervals and visualized using IGV v2.19.7(78). Genes located within centromeric regions were identified using Bedtools v2.31.1(79) based on gene annotations; a gene was assigned to a centromeric region only if more than 50% of its total length overlapped with the centromeric interval.

### DEG identification and enrichment analysis

These clean RNA-seq reads were then aligned to the T2T *D. officinale* genome using HISAT2. Gene expression levels were quantified using featureCounts v2.0.6(80), generating a count matrix for downstream analysis. In addition to raw counts, TPM values were calculated for visualization purposes.

Prior to differential expression analysis, expression data were filtered by retaining only genes with counts ≥ 10 in at least five out of nine samples. PCA and hierarchical clustering were performed on vst-transformed expression data to assess sample relationships and quality.

Differential expression analysis was performed using DESeq2 v1.10 (81), with significance thresholds set at | log_2_(FoldChange)| > 1 and adjusted *p*-value (Padj) < 0.05. Three pairwise comparisons were conducted: stem vs root, stem vs leaf, and leaf vs root. The numbers of differentially expressed genes (DEGs) identified in each comparison were summarized, and their overlaps were visualized using Venn diagrams.

For functional analysis, we focused on identifying tissue-specifically up-regulated genes. Specifically, genes that were significantly up-regulated in both of the two relevant pairwise comparisons (e.g., up in leaf vs. stem and up in leaf vs. root) were defined as genes up-regulated in that tissue (i.e., leaf-upregulated, stem-upregulated, or root-upregulated genes). These up-regulated genes were subjected to GO and KEGG enrichment analyses using clusterProfiler(82). For each enrichment analysis, genes that both passed the expression filter and possessed functional annotation were used as the background gene set. Statistical significance was evaluated using the hypergeometric test with Benjamini–Hochberg correction, and pathways or terms with Padj < 0.05 were considered significantly enriched.

### Gene family analysis

For comparative genomic analyses, protein sequences from the T2T assembly of *D. officinale* and 24 additional plant genomes were collected. Genomic data for *Dendro-bium nobile* (GCA 022539455.1), *Dendrobium chrysotoxum* (GCA 019925795.1), *Dendrobium thyr-siflorum* (GCA 040670175.1), *Dendrobium huoshanense* (GCA 016618105.1), *Phalaenopsis equestris* (GCF 001263595.1), *Gastrodia elata* (GWHBDNU00000000), *Apostasia shenzhenica* (GCA 002786265.1), *Oryza sativa* (http://ricesuperpir.com/web/download), *Brachypodium distachyon* (GCF 000005505.3) and *Arabidopsis thaliana* (GCF 000001735.4). In addition, protein sequences of fourteen *Dendrobium* species, including *Dendrobium chao Praya Smile*, *Dendrobium discolor*, *Dendrobium crocatum*, *Dendrobium smilliae*, *Dendrobium leonis*, *Dendrobium crumenatum*, *Dendrobium formosum*, *Dendrobium secundum*, *Dendrobium ellipsophyllum*, *Dendrobium tetragonum*, *Dendrobium hercoglossum*, *Dendrobium lindleyi*, *Dendrobium aphyllum*, *Dendrobium jenkinsii*, were downloaded from Figshare (https://doi.org/10.6084/m9.figshare.26342338)(83). These sequences were used to infer orthologous genes across these species using OrthoFinder v2.5.5(84).

### Phylogenomic analysis

The resulting single-copy orthologous gene sequences served as the foundation for subsequent analyses. Multiple sequence alignment of these sequences was performed using Muscle v5.1(85), and conserved sites were extracted using Gblocks v0.91b(86; 87) to improve the reliability of the phylogenetic analysis. To select the most appropriate evolutionary model, ModelTest-NG v0.2.0(88) was employed. A phylogenetic tree was then constructed using IQ-TREE v2.3.6(89) based on the maximum likelihood method.

To estimate species divergence times, the MCMCTree module in PAML v4.0 (90) was used. Species divergence time estimates were queried from the TimeTree database(91) and incorporated as calibration constraints in the molecular dating analysis. These calibration constraints included the divergence between *Oryza sativa* and *Brachypodium distachyon* (41.5–62.0 Mya), *Apostasia shenzhenica* and *Oryza sativa* (108.4–125.0 Mya), *Apostasia shenzhenica* and *Gastrodia elata* (72.2–114.1 Mya), as well as the split between *Phalaenopsis equestris* and *Dendrobium* species (12.3–51.0 Mya).

Based on the gene family clustering results, CAFE v5.1(92) was used to infer gene family expansion and contraction across the species phylogeny. Expanded and contracted gene families identified on the *D. officinale* node were extracted and subjected to GO and KEGG enrichment analyses using clusterProfiler. All *D. officinale* genes that were assigned to gene families, included in the CAFE analysis, and possessed GO or KEGG annotations were used as the background gene set.

### Whole-genome duplication analysis

WGD analysis was performed using WGDI (93). Syntenic blocks within the haplotype-resolved *D. officinale* genome were identified, and homologous gene pairs within these blocks were extracted based on sequence similarity and genomic collinearity. The Ks was calculated for paralogous gene pairs within *D. officinale* as well as for orthologous syntenic gene pairs between *D. officinale* and several other species, including *P. equestris*, *V. vinifera* (https://grapedia.org/files-download/), *G. elata*, *D. chrysotoxum* and *D. huoshanense*. Using JCVI(94), synteny was investigated between *D. chrysotoxum* and the four haplotypes of *D. officinale*, as well as between *A. trichopoda* (GWHFSRF00000000.1) and *D. chrysotoxum*. The synonymous substitution rate (*r*) for *D. officinale* was calibrated based on the absolute age of the orchid-shared WGD event (74 Mya)(28) and the corresponding *K_s_*peak value in *D. officinale* (Ks = 0.89). Using these values, *r* was estimated to be 6.01 × 10^−9^ substitutions per site per year. This calibrated rate was then applied in the formula *T* = *K_s_/*(2*r*) to estimate the timing of the recent WGD event in *D. officinale*.

### Identification of the SWEET family in *Dendrobium*

To investigate the *SWEET* gene family in 19 *Dendrobium* species (*D. chao Praya Smile*, *D. discolor*, *D. crocatum*, *D. smilliae*, *D. leonis*, *D. crumenatum*, *D. formosum*, *D. secundum*, *D. ellipsophyllum*, *D. tetragonum*, *D. hercoglossum*, *D. lindleyi*, *D. aphyllum*, *D. jenkinsii*, *D. nobile*, *D. chrysotoxum*, *D. thyrsiflorum*, *D. huoshanense*, and *D. officinale*), candidate genes were first identified using a combination of sequence similarity and domain-based screening. Seventeen SWEET protein sequences from *Arabidopsis thaliana* and 21 from *Oryza sativa* were obtained and used as queries in BLASTP searches against the proteomes of the 19 *Dendrobium* species, with the parameters -outfmt 6 -evalue 1e-5. Subsequently, domain-based screening was performed with HMMER (hmmsearch) using the Pfam HMM profile PF03083 to detect candidates. All candidate genes were subsequently analyzed using InterPro and the NCBI Conserved Domain Database to confirm the presence of conserved domains. Only genes containing at least one complete MtN3/saliva domain were retained as *SWEET* genes (95; 96). Microsynteny analysis of *SWEET* genes among *Dendrobium* species was performed using JCVI.

Multiple sequence alignment of the identified SWEET protein sequences from 19 *Dendrobium* species, together with those from *Arabidopsis thaliana* and *Oryza sativa*, was performed using Muscle. A maximum likelihood (ML) phylogenetic tree was constructed using IQ-TREE with a bootstrap value of 1000. Additionally, physicochemical properties—including amino acid number, pI, molecular weight, instability index, hydrophobicity, and aliphatic index—of the *DoffSWEET* members were analyzed using Expasy tools(97). Subcellular localization predictions for all members were conducted with the CellPLoc2.0 online tool(98).

### Identification of gene duplication modes

Gene duplication modes were identified using Dup-Gen finder v1.0.0(99), which classifies gene pairs into whole-genome duplication, tandem duplication, proximal duplication, transposed duplication, and dispersed duplication. The analysis was based on an all-vs-all BLASTP search with parameters -outfmt 6 -evalue 1e-5.

### Cis-acting elements analysis in *D. officinale*

The 2,000 bp upstream sequences of the start codon of *SWEET* genes were extracted and analyzed for cis-acting regulatory elements using PlantCARE database (https://bioinformatics.psb.ugent.be/webtools/plantcare/html/). Unnamed elements, as well as basic elements such as TATA-box and CAAT-box, were excluded from the analysis. The identified elements were classified into four major categories: Light responsive, Plant growth and development, Phytohormone responsive, and abiotic and biotic stress elements(100; 101).

### Expression analysis of haplotype genomes

Gene structure annotation of the four haplotype genomes was performed using an integrated annotation approach, combining transcriptome-based prediction, homology-based prediction, and ab initio prediction using the same pipeline applied to the T2T reference genome, with the addition of ab initio prediction using ANNEVO v2.1(102). To further improve completeness, T2T annotations were projected onto each haplotype genome using Liftoff v1.6.3(103) to supplement the primary annotations. Subsequently, pairwise comparisons among the four haplotypes were conducted with MCScanX v1.0.0(104) to identify syntenic blocks. Allelic genes were defined as those gene pairs located within syntenic blocks that exhibited a strict one-to-one alignment relationship(105; 106).

A composite reference genome, consisting of the concatenated assemblies of all four haplotypes, was used for mapping cleaned RNA-seq reads. Only uniquely aligned reads were retained for downstream analyses. Gene expression levels were quantified using StringTie and measured in Transcripts Per Million (TPM). Alleles within tetrads with a total TPM > 0.5 across all four copies were defined as “expressed”, following a previously established wheat transcriptome-based approach (107). To allow direct comparisons among alleles within each tetrad, relative expression values were calculated for each allele as TPM of that allele divided by the sum of TPMs across the four alleles (e.g., TPM(A)/[TPM(A) + TPM(B) + TPM(C) + TPM(D)]). Based on the relative expression values, expression patterns were classified into 15 categories, including a balanced pattern and 14 biased patterns. Dominant patterns included single-allele dominance (hapA-, hapB-, hapC-, or hapD-dominant) and dual-allele dominance (hapAhapB-, hapAhapC-, hapAhapD-, hapBhapC-, hapBhapD-, or hapChapD-dominant), while suppressed patterns included single-allele suppression (hapA-, hapB-, hapC-, or hapD-suppress)(105). Finally, Euclidean distance (ED) to each of the 15 idealized expression patterns was computed for each tetrad, and the pattern with the minimum ED was used to assign its expression category.

For functional enrichment analyses, we focused on root tissue, which exhibited the highest number of expressed alleles across all tetrads. The 15 allelic expression categories were consolidated into four major classes. Single-allele dominant expression included tetrads with dominance of a single allele (hapA-, hapB-, hapC-, or hapD-dominant). Dual-allele dominant expression included tetrads with co-dominance of two alleles (hapAhapB-, hapAhapC-, hapAhapD-, hapBhapC-, hapBhapD-, or hapChapD-dominant). Single-allele suppressed expression included tetrads in which one allele showed reduced expression relative to the others (hapA-, hapB-, hapC-, or hapD-suppressed). Balanced expression included tetrads in which all four alleles exhibited similar relative expression. For each tetrad, only the hapA copy was retained as a representative. GO and KEGG enrichment analyses were subsequently performed on these hapA gene sets using the clusterProfiler R package, with hapA copies representing all tetrads expressed in root tissue serving as the background.

### Haplotype collinearity and genomic variation analysis

All-vs-all protein sequence comparisons were first performed using BLASTP (*E*-value < 10^−5^), followed by collinearity analysis with MCScanX. Syntenic relationships among the four haplotypes of *D. officinale* were visualized using ChiPlot (https://www.chiplot.online/McScanX_synteny_plot.html).

Pairwise whole-genome alignments were performed to compare (i) the four haplotype assemblies of *D. officinale* with each other and (ii) each haplotype against the *D. officinale* v3.0 reference genome. Single-nucleotide polymorphisms (SNPs) were identified from the nucmer v4.0.1 alignments (108), whereas insertions/deletions (Indels) and structural variations (SVs) were detected using Assemblytics v1.2.1 (109).

## Acknowledgements

XL is supported by the Natural Science Foundation of Hunan Province (Grant No. 2024JJ4008), the National Natural Science Foundation of China (Grant No. 32400506), and Fundamental Research Funds for the Central Universities (Grant No. 541109030062). XG is supported by the National Natural Science Foundation of China (Grant No. 32372124) and the Key Research & Development Project of Hunan Provincial Department of Science and Technology (2023NK2022). YL is supported by the Yuelushan Laboratory Breeding Program (YLS-2025-ZY03024).

## Author contributions

Xiao Luo, Xinhong Guo, Yibo Luo conceived this study. Enlian Chen, Jialu Xu, Yuansheng Liu, Yiping Feng, Shance Niu and Qinghua Lu conducted the data analysis. Yongliang Li collected the samples. Xiaoyu Ding and Zhitao Niu provided the genome annotation information of *D. officinale* v3.0. Enlian Chen and Jialu Xu drafted the manuscript. Xiao Luo, Yibo Luo, Xinhong Guo, Yuansheng Liu and Si Qin revised the manuscript. All authors read and approved the final version of the manuscript.

## Data Availability

The raw sequencing data—including PacBio, Oxford Nanopore Technologies (ONT), Hi-C, paired-end NGS sequencing, and RNA-seq—have been deposited in the China National Genomics Data Center (https://bigd.big.ac.cn/) under BioProject accession number **PRJCA040583**. The corresponding BioSample accession numbers are **SAMC5130779** through **SAMC5130789**. The T2T genome assembly of *Dendrobium officinale* is available via NCBI under project accession **PRJNA1267708**. Additionally, genome assemblies for the four phased haplotypes (hapA, hapB, hapC, and hapD) have been submitted to NCBI under accession numbers **PRJNA1267690**, **PRJNA1267689**, **PRJNA1267688**, and **PRJNA1267687**, respectively.

## References

[1] Wood, H. P. The dendrobiums (ARG Gantner Verlag, 2006).

[2] Moudi, M., Go, R., Yien, C. Y. S. & Saleh, M. N. A review on molecular systematic of the genus dendrobium sw. Acta Biologica Malaysiana 2, 71–78 (2013).

[3] Chen, X., Chen, C. & Fu, X. Dendrobium officinale polysaccharide alleviates type 2 diabetes mellitus by restoring gut microbiota and repairing intestinal barrier via the lps/tlr4/trif/nf-kb axis. Journal of Agricultural and Food Chemistry 71, 11929–11940 (2023).

[4] Guo, L. et al. Dendrobium officinale kimura & migo polysaccharide and its multilayer emulsion protect skin photoaging. Journal of Ethnopharmacology 318, 116974 (2024).

[5] Hui, A. et al. A comparative study of pectic polysaccharides from fresh and dried dendrobium officinale based on their structural properties and hepatoprotection in alcoholic liver damaged mice. Food & Function 14, 4267–4279 (2023).

[6] Wang, Y. et al. Dendrobium offificinale polysaccharides prevents glucocorticoids-induced osteoporosis by destabilizing keap1-nrf2 interaction. International Journal of Biological Macromolecules 253, 126600 (2023).

[7] Zhang, Y. et al. Dendrobium officinale kimura & migo attenuates high cholesterol diet-induced atherosclerosis. Natural Product Communications 19, 1934578X241282497 (2024).

[8] Schuiteman, A. The dendrobiums (2007).

[9] Yuan, Y., Yu, M., Zhang, B., Liu, X. & Zhang, J. Comparative nutritional characteristics of the three major chinese dendrobium species with different growth years. PLoS One 14, e0222666 (2019).

[10] Yan, L. et al. The genome of dendrobium officinale illuminates the biology of the important traditional chinese orchid herb. Molecular plant 8, 922–934 (2015).

[11] Zhang, G.-Q. et al. The dendrobium catenatum lindl. genome sequence provides insights into polysaccharide synthase, floral development and adaptive evolution. Scientific reports 6, 19029 (2016).

[12] Niu, Z. et al. The chromosome-level reference genome assembly for dendrobium officinale and its utility of functional genomics research and molecular breeding study. Acta Pharmaceutica Sinica B 11, 2080–2092 (2021).

[13] Liu, Y. et al. Integrated metabolomic and transcriptomic analysis reveals the mechanism of high polysaccharide content in tetraploid dendrobium catenatum lindl. Industrial Crops and Products 212, 118391 (2024).

[14] Pham, P.-L. et al. Changes in morphological characteristics, regeneration ability, and polysaccharide content in tetraploid dendrobium officinale. HortScience 54, 1879–1886 (2019).

[15] Yeung, E. C. A perspective on orchid seed and protocorm development. Botanical studies 58, 33 (2017).

[16] Xu, Z.-X. et al. Symbiosis between dendrobium catenatum protocorms and serendipita indica involves the plant hypoxia response pathway. Plant Physiology 192, 2554–2568 (2023).

[17] Zhao, D.-K., Mou, Z.-M. & Ruan, Y.-L. Orchids acquire fungal carbon for seed germination: pathways and players. Trends in Plant Science 29, 733–741 (2024).

[18] Rasmussen, H. N. & Rasmussen, F. N. Orchid mycorrhiza: implications of a mycophagous life style. Oikos 118, 334–345 (2009).

[19] Cheng, H., Concepcion, G. T., Feng, X., Zhang, H. & Li, H. Haplotype-resolved de novo assembly using phased assembly graphs with hifiasm. Nature methods 18, 170–175 (2021).

[20] Li, H. & Durbin, R. Fast and accurate short read alignment with burrows–wheeler transform. bioinformatics 25, 1754–1760 (2009).

[21] Zeng, X. et al. Chromosome-level scaffolding of haplotype-resolved assemblies using hi-c data without reference genomes. Nature Plants 10, 1184–1200 (2024).

[22] Durand, N. C. et al. Juicebox provides a visualization system for hi-c contact maps with unlimited zoom. Cell systems 3, 99–101 (2016).

[23] Xu, M. et al. Tgs-gapcloser: a fast and accurate gap closer for large genomes with low coverage of error-prone long reads. GigaScience 9, giaa094 (2020).

[24] Simao, F. A., Waterhouse, R. M., Ioannidis, P., Kriventseva, E. V. & Zdobnov, E. M. Busco: assessing genome assembly and annotation completeness with single-copy orthologs. Bioinformatics 31, 3210–3212 (2015).

[25] Rhie, A., Walenz, B. P., Koren, S. & Phillippy, A. M. Merqury: reference-free quality, completeness, and phasing assessment for genome assemblies. Genome biology 21, 1–27 (2020).

[26] Benson, G. Tandem repeats finder: a program to analyze dna sequences. Nucleic acids research 27, 573–580 (1999).

[27] Zhang, R. Centromics: A tool for centromere analysis. https://github.com/zhangrengang/Centromics (2023).

[28] Zhang, G.-Q. et al. The apostasia genome and the evolution of orchids. Nature 549, 379–383 (2017).

[29] Ming, R. et al. The pineapple genome and the evolution of cam photosynthesis. Nature genetics 47, 1435–1442 (2015).

[30] Zhang, Y. et al. Chromosome-scale assembly of the dendrobium chrysotoxum genome enhances the understanding of orchid evolution. Horticulture Research 8 (2021).

[31] Xu, Q. et al. Chromosome-scale assembly of the dendrobium nobile genome provides insights into the molecular mechanism of the biosynthesis of the medicinal active ingredient of dendrobium. Frontiers in genetics 13, 844622 (2022).

[32] Eom, J.-S. et al. Sweets, transporters for intracellular and intercellular sugar translocation. Current opinion in plant biology 25, 53–62 (2015).

[33] Durand, M. et al. Carbon source–sink relationship in arabidopsis thaliana: the role of sucrose transporters. Planta 247, 587–611 (2018).

[34] Valifard, M. et al. Vacuolar fructose transporter sweet17 is critical for root development and drought tolerance. Plant Physiology 187, 2716–2730 (2021).

[35] Wei, Y., Xiao, D., Zhang, C. & Hou, X. The expanded sweet gene family following whole genome triplication in brassica rapa. Genes 10, 722 (2019).

[36] Jiang, Y.-j., Peng, Z., Yu, W., et al. Genome-wide identification and transcriptome profiling reveal great expansion of sweet gene family and their wide-spread responses to abiotic stress in wheat (triticum aestivum l.). Journal of Integrative Agriculture 19, 1704–1720 (2020).

[37] Chardon, F. et al. Leaf fructose content is controlled by the vacuolar transporter sweet17 in arabidopsis. Current Biology 23, 697–702 (2013).

[38] Klemens, P. A. et al. Overexpression of the vacuolar sugar carrier atsweet16 modifies germination, growth, and stress tolerance in arabidopsis. Plant Physiology 163, 1338–1352 (2013).

[39] Seo, P. J., Park, J.-M., Kang, S. K., Kim, S.-G. & Park, C.-M. An arabidopsis senescence-associated protein sag29 regulates cell viability under high salinity. Planta 233, 189–200 (2011).

[40] Julius, B. T., Leach, K. A., Tran, T. M., Mertz, R. A. & Braun, D. M. Sugar transporters in plants: new insights and discoveries. Plant and Cell Physiology 58, 1442–1460 (2017).

[41] Durand, M. et al. Water deficit enhances c export to the roots in arabidopsis thaliana plants with contribution of sucrose transporters in both shoot and roots. Plant physiology 170, 1460–1479 (2016).

[42] Zheng, L. et al. The soybean sugar transporter gmsweet6 participates in sucrose transport towards fungi during arbuscular mycorrhizal symbiosis. Plant, Cell & Environment 47, 1041–1052 (2024).

[43] An, J. et al. A medicago truncatula sweet transporter implicated in arbuscule maintenance during arbuscular mycorrhizal symbiosis. New Phytologist 224, 396–408 (2019).

[44] Li, L. et al. The sweet14 sugar transporter mediates mycorrhizal symbiosis and carbon allocation in dendrobium officinale. BMC Plant Biology 25, 416 (2025).

[45] Waterman, R. J. & Bidartondo, M. I. Deception above, deception below: linking pollination and mycorrhizal biology of orchids. Journal of Experimental Botany 59, 1085–1096 (2008).

[46] Suetsugu, K. & Matsubayashi, J. Evidence for mycorrhizal cheating in apostasia nipponica, an early-diverging member of the orchidaceae. New Phytologist 229, 2302–2310 (2021).

[47] Chen, S., Zhou, Y., Chen, Y. & Gu, J. fastp: an ultra-fast all-in-one fastq preprocessor. Bioinformatics 34, i884–i890 (2018).

[48] Marcais, G. & Kingsford, C. A fast, lock-free approach for efficient parallel counting of occurrences of k-mers. Bioinformatics 27, 764–770 (2011).

[49] Ranallo-Benavidez, T. R., Jaron, K. S. & Schatz, M. C. Genomescope 2.0 and smudgeplot for reference-free profiling of polyploid genomes. Nature communications 11, 1432 (2020).

[50] Guan, D. et al. Identifying and removing haplotypic duplication in primary genome assemblies. Bioinformatics 36, 2896–2898 (2020).

[51] Hu, J., Fan, J., Sun, Z. & Liu, S. Nextpolish: a fast and efficient genome polishing tool for long-read assembly. Bioinformatics 36, 2253–2255 (2020).

[52] Jain, C., Rhie, A., Hansen, N. F., Koren, S. & Phillippy, A. M. Long-read mapping to repetitive reference sequences using winnowmap2. Nature methods 19, 705–710 (2022).

[53] Li, H. Minimap2: pairwise alignment for nucleotide sequences. Bioinformatics 34, 3094–3100 (2018).

[54] Hu, J. et al. Nextpolish2: a repeat-aware polishing tool for genomes assembled using hifi long reads. Genomics, Proteomics & Bioinformatics 22, qzad009 (2024).

[55] Garrison, E. & Marth, G. Haplotype-based variant detection from short-read sequencing. arXiv preprint arXiv:1207.3907 (2012).

[56] Schrinner, S. D. et al. Haplotype threading: accurate polyploid phasing from long reads. Genome biology 21, 1–22 (2020).

[57] Xu, Z. & Wang, H. Ltr_finder: an efficient tool for the prediction of full-length ltr retrotransposons. Nucleic acids research 35, W265–W268 (2007).

[58] Ellinghaus, D., Kurtz, S. & Willhoeft, U. Ltrharvest, an efficient and flexible software for de novo detection of ltr retrotransposons. BMC bioinformatics 9, 1–14 (2008).

[59] Ou, S. & Jiang, N. Ltr_retriever: a highly accurate and sensitive program for identification of long terminal repeat retrotransposons. Plant physiology 176, 1410–1422 (2018).

[60] Flynn, J. M. et al. Repeatmodeler2 for automated genomic discovery of transposable element families. Proceedings of the National Academy of Sciences 117, 9451–9457 (2020).

[61] Chen, N. Using repeat masker to identify repetitive elements in genomic sequences. Current protocols in bioinformatics 5, 4–10 (2004).

[62] Chen, J. et al. A complete telomere-to-telomere assembly of the maize genome. Nature genetics 55, 1221–1231 (2023).

[63] Kim, D., Paggi, J. M., Park, C., Bennett, C. & Salzberg, S. L. Graph-based genome alignment and genotyping with hisat2 and hisat-genotype. Nature biotechnology 37, 907–915 (2019).

[64] De Coster, W., D’hert, S., Schultz, D. T., Cruts, M. & Van Broeckhoven, C. Nanopack: visualizing and processing long-read sequencing data. Bioinformatics 34, 2666–2669 (2018).

[65] epi2me-labs. Pychopper: Full-length cdna read identification and trimming tool. https://github.com/epi2me-labs/pychopper (2022).

[66] Shumate, A., Wong, B., Pertea, G. & Pertea, M. Improved transcriptome assembly using a hybrid of long and short reads with stringtie. PLoS computational biology 18, e1009730 (2022).

[67] Haas, B. J. Transdecoder: Identify candidate coding regions within transcript sequences. https://github.com/TransDecoder/TransDecoder (2023). Software version: [5.7.1].

[68] Li, H. Protein-to-genome alignment with miniprot. Bioinformatics 39, btad014 (2023).

[69] Stanke, M., Tzvetkova, A. & Morgenstern, B. Augustus at egasp: using est, protein and genomic alignments for improved gene prediction in the human genome. Genome biology 7, 1–8 (2006).

[70] Holst, F. et al. Helixer–de novo prediction of primary eukaryotic gene models combining deep learning and a hidden markov model. BioRxiv 2023–02 (2023).

[71] Lomsadze, A., Ter-Hovhannisyan, V., Chernoff, Y. O. & Borodovsky, M. Gene identification in novel eukaryotic genomes by self-training algorithm. Nucleic acids research 33, 6494–6506 (2005).

[72] Haas, B. J. et al. Automated eukaryotic gene structure annotation using evidencemodeler and the program to assemble spliced alignments. Genome biology 9, 1–22 (2008).

[73] Jones, P. et al. Interproscan 5: genome-scale protein function classification. Bioinformatics 30, 1236–1240 (2014).

[74] Seemann, T. Barrnap: Basic rapid ribosomal rna predictor. https://github.com/tseemann/barrnap (2018).

[75] Chan, P. P., Lin, B. Y., Mak, A. J. & Lowe, T. M. trnascan-se 2.0: improved detection and functional classification of transfer rna genes. Nucleic acids research 49, 9077–9096 (2021).

[76] Nawrocki, E. P. & Eddy, S. R. Infernal 1.1: 100-fold faster rna homology searches. Bioinformatics 29, 2933–2935 (2013).

[77] Shen, T. et al. Haplotype-resolved telomere-to-telomere genome assembly of populus lasiocarpa unveils retrotransposon-driven centromere evolution. The Plant Journal 123, e70504 (2025).

[78] Robinson, J. T. et al. Integrative genomics viewer. Nature biotechnology 29, 24–26 (2011).

[79] Quinlan, A. R. & Hall, I. M. Bedtools: a flexible suite of utilities for comparing genomic features. Bioinformatics 26, 841–842 (2010).

[80] Liao, Y., Smyth, G. K. & Shi, W. featurecounts: an efficient general purpose program for assigning sequence reads to genomic features. Bioinformatics 30, 923–930 (2014).

[81] Love, M. I., Huber, W. & Anders, S. Moderated estimation of fold change and dispersion for rna-seq data with deseq2. Genome biology 15, 1–21 (2014).

[82] Wu, T. et al. clusterprofiler 4.0: A universal enrichment tool for interpreting omics data. The innovation 2 (2021).

[83] Li, Y. et al. Pangeneric genome analyses reveal the evolution and diversity of the orchid genus dendrobium. Nature Plants 11, 421–437 (2025).

[84] Emms, D. M. & Kelly, S. Orthofinder: phylogenetic orthology inference for comparative genomics. Genome biology 20, 1–14 (2019).

[85] Edgar, R. C. Muscle5: High-accuracy alignment ensembles enable unbiased assessments of sequence homology and phylogeny. Nature Communications 13, 6968 (2022).

[86] Talavera, G. & Castresana, J. Improvement of phylogenies after removing divergent and ambiguously aligned blocks from protein sequence alignments. Systematic biology 56, 564–577 (2007).

[87] Castresana, J. Selection of conserved blocks from multiple alignments for their use in phylogenetic analysis. Molecular biology and evolution 17, 540–552 (2000).

[88] Darriba, D. et al. Modeltest-ng: a new and scalable tool for the selection of dna and protein evolutionary models. Molecular biology and evolution 37, 291–294 (2020).

[89] Minh, B. Q. et al. Iq-tree 2: new models and efficient methods for phylogenetic inference in the genomic era. Molecular biology and evolution 37, 1530–1534 (2020).

[90] Yang, Z. Paml 4: phylogenetic analysis by maximum likelihood. Molecular biology and evolution 24, 1586–1591 (2007).

[91] Kumar, S. et al. Timetree 5: an expanded resource for species divergence times. Molecular biology and evolution 39, msac174 (2022).

[92] Mendes, F. K., Vanderpool, D., Fulton, B. & Hahn, M. W. Cafe 5 models variation in evolutionary rates among gene families. Bioinformatics 36, 5516–5518 (2020).

[93] Sun, P. et al. Wgdi: a user-friendly toolkit for evolutionary analyses of whole-genome duplications and ancestral karyotypes. Molecular plant 15, 1841–1851 (2022).

[94] Tang, H., et al. Jcvi: A versatile toolkit for comparative genomics analysis. Imeta 3, e211 (2024).

[95] Xuan, Y. H. et al. Functional role of oligomerization for bacterial and plant sweet sugar transporter family. Proceedings of the National Academy of Sciences 110, E3685–E3694 (2013).

[96] Yuan, M. & Wang, S. Rice mtn3/saliva/sweet family genes and their homologs in cellular organisms. Molecular plant 6, 665–674 (2013).

[97] Gasteiger, E. et al. Expasy: the proteomics server for in-depth protein knowledge and analysis. Nucleic acids research 31, 3784–3788 (2003).

[98] Chou, K.-C. & Shen, H.-B. Cell-ploc 2.0: an improved package of web-servers for predicting subcellular localization of proteins in various organisms. Natural Science 2, 1090–1103 (2010).

[99] Qiao, X. et al. Gene duplication and evolution in recurring polyploidization–diploidization cycles in plants. Genome biology 20, 38 (2019).

[100] Wang, Y. et al. Genome-wide identification, expression profiling, and protein interaction analysis of the ccoaomt gene family in the tea plant (camellia sinensis). BMC genomics 25, 238 (2024).

[101] Mengarelli, D. A. & Zanor, M. I. Genome-wide characterization and analysis of the cct motif family genes in soybean (glycine max). Planta 253, 15 (2021).

[102] Ye, K., et al. Highly accurate ab initio gene annotation with annevo (2025).

[103] Shumate, A. & Salzberg, S. L. Liftoff: accurate mapping of gene annotations. Bioinformatics 37, 1639–1643 (2021).

[104] Wang, Y. et al. Mcscanx: a toolkit for detection and evolutionary analysis of gene synteny and collinearity. Nucleic acids research 40, e49–e49 (2012).

[105] Bao, Z. et al. Genome architecture and tetrasomic inheritance of autotetraploid potato. Molecular plant 15, 1211–1226 (2022).

[106] Han, X. et al. Two haplotype-resolved, gap-free genome assemblies for actinidia latifolia and actinidia chinensis shed light on the regulatory mechanisms of vitamin c and sucrose metabolism in kiwifruit. Molecular Plant 16, 452–470 (2023).

[107] Ramırez-Gonzalez, R., et al. The transcriptional landscape of polyploid wheat. Science 361, eaar6089 (2018).

[108] Marcais, G., et al. Mummer4: A fast and versatile genome alignment system. PLoS computational biology 14, e1005944 (2018).

[109] Nattestad, M. & Schatz, M. C. Assemblytics: a web analytics tool for the detection of variants from an assembly. Bioinformatics 32, 3021–3023 (2016).

