## Supplementary File 1 for "Telomere-to-telomere assembly and haplotype analysis of tetraploid *Dendrobium officinale* illuminate Orchidaceae polyploid evolution and mycorrhizal symbiosis genes"

### Supplementary Tables and Figures

**Supplementary Table 1.** Summary of sequencing data used in this study. The sequencing depth was calculated based on a haploid genome size. Q20 represents the percentage of bases with Phred quality score $\geq 20$.

| Types | Clean base (Gbp) | Depth (X) | Q20 (%) | Sample |
|---|---|---|---|---|
| DNBSEQ | 120.0 | ~120 | 98.31 | leaf |
| HiFi | 167.1 | ~167 | 84.68 | leaf |
| ONT | 168.4 | ~168 | 51.61 | leaf |
| ulONT | 67.7 | ~67 | 51.59 | leaf |
| Hi-C | 226.0 | ~226 | 97.53 | leaf |
| RNA-seq (NGS) | 14.7 | - | 97.99 | leaf_1 |
| RNA-seq (NGS) | 12.9 | - | 97.94 | leaf_2 |
| RNA-seq (NGS) | 13.9 | - | 97.92 | leaf_3 |
| RNA-seq (NGS) | 14.5 | - | 98.03 | stem_1 |
| RNA-seq (NGS) | 14.3 | - | 98.09 | stem_2 |
| RNA-seq (NGS) | 12.6 | - | 97.98 | stem_3 |
| RNA-seq (NGS) | 12.7 | - | 97.91 | root_1 |
| RNA-seq (NGS) | 13.9 | - | 97.85 | root_2 |
| RNA-seq (NGS) | 10.5 | - | 98.00 | root_3 |
| RNA-seq (ONT) | 17.5 | - | 61.88 | leaf |

**Supplementary Table 2.** Telomeric positions in *D. officinale* T2T assembly.

| Chromosome | Left Start (bp) | Left End (bp) | Right Start (bp) | Right End (bp) |
|---|---|---|---|---|
| chr1 | 1 | 1,883 | 87,755,520 | 87,770,857 |
| chr2 | 1 | 51,870 | 77,694,476 | 77,696,338 |
| chr3 | 1 | 5,698 | 77,615,009 | 77,618,992 |
| chr4 | 1 | 2,758 | 81,394,072 | 81,396,214 |
| chr5 | 1 | 2,436 | 80,010,067 | 80,012,377 |
| chr6 | 1 | 1,911 | 82,141,964 | 82,143,616 |
| chr7 | 1 | 6,216 | 77,127,411 | 77,128,846 |
| chr8 | 1 | 8,218 | 61,668,372 | 61,670,248 |
| chr9 | 1 | 5,635 | 55,243,229 | 55,249,851 |
| chr10 | 1 | 5,523 | 59,648,473 | 59,679,217 |
| chr11 | 1 | 79,618 | 53,567,316 | 53,578,649 |
| chr12 | 1 | 4,067 | 49,978,783 | 49,984,187 |
| chr13 | 1 | 57,764 | 51,779,983 | 51,781,453 |
| chr14 | 1 | 4,774 | 46,782,911 | 46,870,803 |
| chr15 | 1 | 1,876 | 44,199,379 | 44,205,602 |
| chr16 | 1 | 728 | 45,655,369 | 45,722,303 |
| chr17 | 1 | 4,984 | 40,557,363 | 40,561,087 |
| chr18 | 1 | 2,611 | 38,623,066 | 38,624,683 |
| chr19 | 1 | 2,527 | 36,778,924 | 36,780,702 |

**Supplementary Table 3.** Classification of repetitive sequences in the genome.

| Classification | Length (bp) | Percentage (%) |
|---|---|---|
| **Class I: Retrotransposon** | 555,782,322 | 48.39 |
| SINE | 1,661,602 | 0.14 |
| LINE | 73,293,377 | 6.38 |
| LTR-Retrotransposon | 480,827,343 | 41.87 |
| LTR/Copia | 87,210,069 | 7.59 |
| LTR/Gypsy | 364,052,112 | 31.70 |
| Other | 29,565,162 | 2.57 |
| **Class II: DNA Transposon** | 180,971,341 | 15.76 |
| hobo-Activator | 70,044,337 | 6.10 |
| Tc1-IS630-Pogo | 41,830,244 | 3.64 |
| Harbinger | 28,986,305 | 2.52 |
| Other | 40,110,455 | 3.49 |
| Satellites | 3,741,689 | 0.33 |
| Satellite unoverlapped with TEs | 281,209 | 0.02 |
| Minisatellites | 17,576,556 | 1.53 |
| Minisatellites unoverlapped with TEs | 2,515,873 | 0.22 |
| Microsatellites | 7,191,744 | 0.63 |
| Microsatellites unoverlapped with TEs | 5,606,991 | 0.49 |
| Rolling-circles | 1,315,298 | 0.11 |
| Unclassified | 98,211,288 | 8.55 |
| Simple Repeats | 10,524,394 | 0.92 |
| **Total Repeats** | **855,208,716** | **74.46** |

**Supplementary Table 4.** Gene structural statistics based on different annotation strategies.

| Gene set | Tools/species | Gene number | Average gene length (bp) | Average CDS length (bp) | Average exon number per gene | Average exon length (bp) | Average intron length (bp) |
|---|---|---|---|---|---|---|---|
| RNAseq | TransDecoder | 26,701 | 15,721.66 | 1,220.17 | 4.71 | 486.86 | 3,254.16 |
| | *D. nobile* | 38,815 | 9,817.99 | 1,233.34 | 3.69 | 334.38 | 3,193.19 |
| | *D. chrysotoxum* | 23,605 | 11,439.94 | 1,084.92 | 4.22 | 256.81 | 3,211.20 |
| Homolog | *D. catenatum* | 38,484 | 7,395.46 | 955.07 | 3.16 | 302.64 | 2,987.54 |
| | *P. equestris* | 31,890 | 16,582.78 | 1,295.40 | 5.85 | 221.39 | 3,151.27 |
| | *D. thyrsiflorum* | 23,986 | 16,508.78 | 1,490.58 | 5.02 | 297.12 | 3,738.88 |
| | Augustus | 40,259 | 10,319.03 | 1,001.97 | 4.10 | 244.35 | 3,005.00 |
| de novo | Helixer | 27,991 | 8,891.04 | 1,084.75 | 5.91 | 250.68 | 1,509.07 |
| | GeneMark | 35,561 | 13,591.69 | 1,125.65 | 5.00 | 225.15 | 3,116.84 |
| Integrated | EVM | 33,457 | 8,409.75 | 1,014.14 | 4.05 | 250.33 | 2,423.88 |
| | PASA | 32,505 | 11,194.26 | 1,059.62 | 4.42 | 309.06 | 2,822.95 |
| | Final set | 30,718 | 11,860.08 | 1,090.21 | 4.57 | 323.95 | 2,877.12 |

**Supplementary Table 5.** Functional annotation of predicted genes for T2T *D. officinale* genome.

| Database | Count | Percentage(%) |
|---|---|---|
| Nr | 28,785 | 93.7 |
| UniProt | 28,573 | 93.0 |
| InterPro | 25,926 | 84.4 |
| GO | 12,939 | 42.1 |
| KEGG | 11,624 | 37.8 |
| Annotated | 29,412 | 95.7 |
| Unannotated | 1,306 | 4.3 |
| Total | 30,718 | 100.0 |

**Supplementary Table 6.** Statistics of non-coding RNA annotation in T2T *D. officinale* genome assembly.

| Type | Copy Number | Total Length (bp) | Average Length (bp) | Percentage (%) |
|---|---|---|---|---|
| miRNA | 87 | 11,055 | 127.07 | 0.00096 |
| tRNA | 450 | 34,150 | 75.89 | 0.00297 |
| rRNA | 4,744 | 1,330,893 | 280.54 | 0.11588 |
| snRNA | 225 | 27,314 | 121.40 | 0.00238 |

**Supplementary Table 7.** Gene and transcript counts for each hapotype.

| Hapotype | genome size | Gene Count | Transcript Count |
|---|---|---|---|
| hapA | 1,030,721,977 | 25,296 | 38,550 |
| hapB | 1,009,236,529 | 25,395 | 36,512 |
| hapC | 1,015,477,179 | 26,110 | 37,946 |
| hapD | 1,019,989,560 | 26,018 | 38,248 |

**Supplementary Table 8.** BUSCO assessment results for genome assembly and annotation.

| | | T2T | hapA | hapB | hapC | hapD |
|---|---|---|---|---|---|---|
| Genome assembly | Complete BUSCOs | 96.8% | 90.8% | 91.0% | 90.3% | 90.8% |
| | Complete and single-copy BUSCOs | 93.5% | 84.9% | 84.6% | 84.3% | 84.8% |
| | Complete Duplicated BUSCOs | 3.3% | 5.9% | 6.4% | 5.9% | 6.0% |
| | Fragmented BUSCOs | 1.1% | 3.4% | 3.7% | 3.5% | 3.7% |
| | Missing BUSCOs | 2.1% | 5.8% | 5.3% | 6.3% | 5.5% |
| | Total BUSCO groups searched | 1614 | 1614 | 1614 | 1614 | 1614 |
| Genome annotation | Complete BUSCOs | 95.0% | 88.0% | 87.4% | 86.8% | 87.9% |
| | Complete and single-copy BUSCOs | 90.80% | 83.1% | 81.3% | 81.5% | 83.5% |
| | Complete Duplicated BUSCOs | 4.2% | 4.8% | 6.1% | 5.3% | 4.4% |
| | Fragmented BUSCOs | 1.4% | 3.2% | 3.8% | 3.1% | 2.4% |
| | Missing BUSCOs | 3.7% | 8.9% | 8.8% | 10.1% | 9.7% |
| | Total BUSCO groups searched | 1614 | 1614 | 1614 | 1614 | 1614 |

**Supplementary Table 9.** Candidate centromeric regions. Centromeres on chr10, chr12, chr14 and chr16 were not found. The centromeric regions on chr3, chr8, chr9, and chr13 remain unverified.

| chromosome | start | end | length | status |
|---|---|---|---|---|
| chr1 | 54,526,532 | 54,662,694 | 136,162 | |
| chr2 | 77,359,040 | 77,662,926 | 303,886 | |
| chr3 | 47,791,388 | 48,127,461 | 336,073 | Unverified |
| chr4 | 26,468,521 | 26,596,157 | 127,636 | |
| chr5 | 25,638,432 | 25,810,780 | 172,348 | |
| chr6 | 40,507,643 | 40,686,806 | 179,163 | |
| chr7 | 76,300,000 | 77,085,977 | 785,977 | |
| chr8 | 34,087,215 | 35,382,566 | 1,295,351 | Unverified |
| chr9 | 54,763,333 | 55,243,069 | 479,736 | Unverified |
| chr10 | – | – | – | – |
| chr11 | 35,880,621 | 35,982,576 | 101,955 | |
| chr12 | – | – | – | – |
| chr13 | 38,905,767 | 39,001,701 | 95,934 | Unverified |
| chr14 | – | – | – | – |
| chr15 | 39,394,789 | 39,517,144 | 122,355 | |
| chr16 | – | – | – | – |
| chr17 | 27,548,120 | 27,640,280 | 92,160 | |
| chr18 | 30,400,000 | 32,416,000 | 2,016,000 | |
| chr19 | 25,652,662 | 26,839,825 | 1,187,163 | |

**Supplementary Table 14.** Variation statistics of *D. officinale*. The table summarizes pairwise comparisons among the four haplotypes and between the *D. officinale* v3.0 reference genome and each haplotype, including the numbers of SNPs, indels, and structural variants (SVs; $> 50$ bp), as well as the total sizes of indels and SVs.

| Haplotype Pair | No. of SNPs | No. of Indels | Indel Size (Mbp) | No. of SVs (>50 bp) | SV Size (Mbp) |
|---|---|---|---|---|---|
| hapA–hapB | 6,300,439 | 1,303,678 | 3.32 | 24,525 | 46.10 |
| hapA–hapC | 6,357,982 | 1,324,867 | 3.36 | 24,910 | 47.03 |
| hapA–hapD | 6,322,502 | 1,315,536 | 3.36 | 24,826 | 46.85 |
| hapB–hapC | 6,283,093 | 1,324,058 | 3.35 | 24,970 | 47.16 |
| hapB–hapD | 6,308,658 | 1,337,717 | 3.40 | 24,780 | 45.90 |
| hapC–hapD | 6,313,992 | 1,332,686 | 3.40 | 25,186 | 47.48 |
| *D. officinale* v3.0–hapA | 7,822,058 | 1,314,320 | 3.94 | 32,297 | 55.85 |
| *D. officinale* v3.0–hapB | 7,739,117 | 1,280,682 | 3.87 | 32,213 | 55.78 |
| *D. officinale* v3.0–hapC | 7,749,904 | 1,291,779 | 3.88 | 32,329 | 54.76 |
| *D. officinale* v3.0–hapD | 7,747,976 | 1,306,536 | 3.92 | 32,359 | 56.26 |

**Supplementary Table 16.**SWEET gene numbers in 19 *Dendrobium* species and associated morphological and ecological traits.

| Accession | Section | SWEET genes | Epiphytic type | Plant size | Geographic distribution | Climate type |
|---|---|---|---|---|---|---|
| D. lindleyi | Densiflora | 15 | Tree | Tiny | South China and Himalaya | Tropical |
| D. ellipsophyllum | Distichophyllae | 16 | Tree | Small | Yunnan, Burma, Lao, Vietnam, Thailand | Tropical |
| D. jenkinsii | Densiflora | 20 | Tree | Tiny | Yunnan, Himalaya | Tropical |
| D. tetragonum | Dendrocoryne | 20 | Rock | Large | Australia | Tropical |
| D. formosum | Formosae | 21 | Tree | Large | Burma, Thailand, India, Nepal, Vietnam | Tropical |
| D. huoshanense | Dendrobium | 21 | Rock | Small | Anhui, Jiangxi, Hunan | Subtropical |
| D. secundum | Pedilonum | 22 | Tree | Large | Burma, Thailand, Vietnam, Malesia, Philippines | Tropical |
| D. discolor | Spatulata | 22 | Tree | Very Large | Australia and New Guinea | Tropical |
| D. thyrsiflorum | Densiflora | 22 | Tree or Rock | Large | Yunnan, Indochina | Tropical |
| D. crumenatum | Crumenata | 24 | Tree or Rock | Small | Sri Lanka, Burma, Thailand, New Guinea, Philippines, Taiwan | Tropical |
| D. leonis | Aporum | 25 | Tree or Rock | Small | Asia and Malesia east to New Guinea | Tropical |
| D. chrysotoxum | Densiflora | 27 | Tree or Rock | Large | Yunnan, Indochina | Tropical |
| D. crocatum | Calcarifera | 28 | Tree or Rock | Large | Malesia | Tropical |
| D. smilliae | Calyptrochilus | 29 | Tree or Rock | Large | New Guinea, Australia | Tropical |
| D. hercoglossum | Breviflores | 30 | Tree or Rock | Large | South China to Thailand, Malesia, Philippines | Tropical |
| D. officinale | Dendrobium | 31 | Tree or Rock | Large | Middle China | Subtropical |
| D. nobile | Dendrobium | 34 | Tree or Rock | Large | Himalaya | Tropical and subtropical |
| D. aphylium | Dendrobium | 40 | Tree or Rock | Large | From Himalaya to Malesia | Tropical |
| D. Chao Praya Smile | Hybrid | 20 | - | - | - | - |

**Supplementary Table 17.**Physicochemical properties of *DoffSWEET* genes in *D. officinale.* Abbreviations: GRAVY, Grand Average of Hydropathicity; Instab. Indx, Instability Index; TMs, transmembrane domains; MtN3, MtN3/saliva domain.

| **Gene Name** | **Protein ID** | **length (aa)** | **Molecular Weight (Da)** | **Theoretical pI** | **Instab. Indx** | **Aliphatic Index** | **GRAVY** | **Subcellular Localization** | **Number of TMs** | **Number of MtN3** |
|---|---|---|---|---|---|---|---|---|---|---|
| DoffSWEET1 | Dof03G000996.mRNA1 | 271 | 30,534.37 | 9.25 | 33.31 | 117.23 | 0.599 | Cell membrane | 7 | 2 |
| DoffSWEET2 | Dof03G001882.mRNA1 | 265 | 29,548.47 | 9.27 | 33.22 | 120.98 | 0.826 | Cell membrane | 7 | 2 |
| DoffSWEET3 | Dof03G001883.mRNA1 | 265 | 29,628.48 | 9.00 | 30.75 | 119.85 | 0.808 | Cell membrane | 7 | 2 |
| DoffSWEET4 | Dof03G002091.mRNA1 | 253 | 27,882.02 | 5.35 | 35.40 | 123.60 | 0.658 | Cell membrane | 7 | 2 |
| DoffSWEET5 | Dof04G000542.mRNA1 | 235 | 26,252.67 | 9.03 | 39.23 | 125.91 | 0.823 | Cell membrane | 7 | 2 |
| DoffSWEET6 | Dof04G000543.mRNA1 | 244 | 27,212.78 | 8.87 | 39.59 | 124.80 | 0.841 | Cell membrane | 7 | 2 |
| DoffSWEET7 | Dof04G000893.mRNA1 | 236 | 26,380.58 | 9.22 | 39.04 | 118.39 | 0.744 | Cell membrane | 7 | 2 |
| DoffSWEET8 | Dof04G001938.mRNA1 | 266 | 29,937.55 | 8.87 | 37.28 | 115.34 | 0.641 | Cell membrane | 7 | 2 |
| DoffSWEET9 | Dof05G000489.mRNA1 | 234 | 26,003.89 | 9.25 | 31.41 | 116.67 | 0.833 | Cell membrane | 7 | 2 |
| DoffSWEET10 | Dof07G001227.mRNA1 | 252 | 28,161.78 | 8.84 | 34.21 | 132.66 | 0.808 | Cell membrane | 7 | 2 |
| DoffSWEET11 | Dof07G001240.mRNA1 | 252 | 28,360.96 | 9.12 | 42.61 | 131.51 | 0.781 | Cell membrane | 7 | 2 |
| DoffSWEET12 | Dof07G001513.mRNA1 | 233 | 26,053.74 | 9.30 | 34.28 | 114.68 | 0.632 | Cell membrane | 7 | 2 |
| DoffSWEET13 | Dof08G000290.mRNA1 | 258 | 29,457.31 | 8.90 | 43.77 | 135.54 | 0.835 | Chloroplast | 7 | 2 |
| DoffSWEET14 | Dof08G001053.mRNA1 | 256 | 28,201.47 | 9.06 | 32.23 | 110.86 | 0.566 | Cell membrane | 7 | 2 |
| DoffSWEET15 | Dof08G001230.mRNA1 | 296 | 33,119.08 | 9.52 | 39.44 | 109.29 | 0.333 | Cell membrane | 7 | 2 |
| DoffSWEET16 | Dof08G000289.mRNA1 | 204 | 23,383.16 | 9.08 | 42.86 | 136.62 | 0.809 | Chloroplast | 5 | 2 |
| DoffSWEET17 | Dof09G000510.mRNA1 | 262 | 29,221.23 | 9.49 | 31.04 | 120.57 | 0.708 | Cell membrane | 7 | 2 |
| DoffSWEET18 | Dof09G000548.mRNA1 | 276 | 30,977.53 | 6.83 | 42.70 | 120.11 | 0.655 | Cell membrane | 7 | 2 |
| DoffSWEET19 | Dof10G000351.mRNA1 | 237 | 26,596.77 | 8.86 | 38.59 | 120.93 | 0.952 | Cell membrane | 7 | 2 |
| DoffSWEET20 | Dof11G000700.mRNA1 | 248 | 27,418.85 | 9.63 | 38.73 | 128.06 | 0.695 | Cell membrane | 7 | 2 |
| DoffSWEET21 | Dof11G000767.mRNA1 | 237 | 26,577.42 | 8.99 | 48.56 | 118.82 | 0.836 | Cell membrane | 7 | 2 |
| DoffSWEET22 | Dof11G000907.mRNA1 | 249 | 27,632.41 | 9.13 | 33.94 | 140.00 | 1.040 | Cell membrane | 7 | 2 |
| DoffSWEET23 | Dof11G001438.mRNA1 | 232 | 25,978.15 | 5.57 | 47.59 | 130.13 | 0.969 | Cell membrane | 7 | 2 |
| DoffSWEET24 | Dof11G001449.mRNA2 | 234 | 26,972.83 | 9.20 | 45.85 | 134.87 | 0.921 | Cell membrane | 7 | 2 |
| DoffSWEET25 | Dof11G001451.mRNA1 | 121 | 13,898.91 | 8.93 | 44.67 | 139.34 | 0.848 | Cell membrane | 3 | 1 |
| DoffSWEET26 | Dof15G000738.mRNA1 | 221 | 25,038.94 | 6.29 | 37.29 | 134.48 | 0.868 | Chloroplast | 6 | 2 |
| DoffSWEET27 | Dof18G000481.mRNA1 | 259 | 29,002.73 | 8.69 | 32.79 | 130.93 | 0.775 | Cell membrane | 7 | 2 |
| DoffSWEET28 | Dof18G000482.mRNA1 | 259 | 28,839.73 | 8.85 | 35.60 | 136.14 | 0.859 | Cell membrane | 7 | 2 |
| DoffSWEET29 | Dof18G000484.mRNA1 | 260 | 28,790.83 | 9.11 | 34.58 | 138.96 | 0.908 | Cell membrane | 7 | 2 |
| DoffSWEET30 | Dof18G000485.mRNA1 | 259 | 28,917.85 | 8.99 | 32.87 | 135.02 | 0.814 | Cell membrane | 7 | 2 |
| DoffSWEET31 | Dof18G000486.mRNA1 | 259 | 28,950.81 | 8.69 | 34.76 | 136.10 | 0.840 | Cell membrane | 7 | 2 |

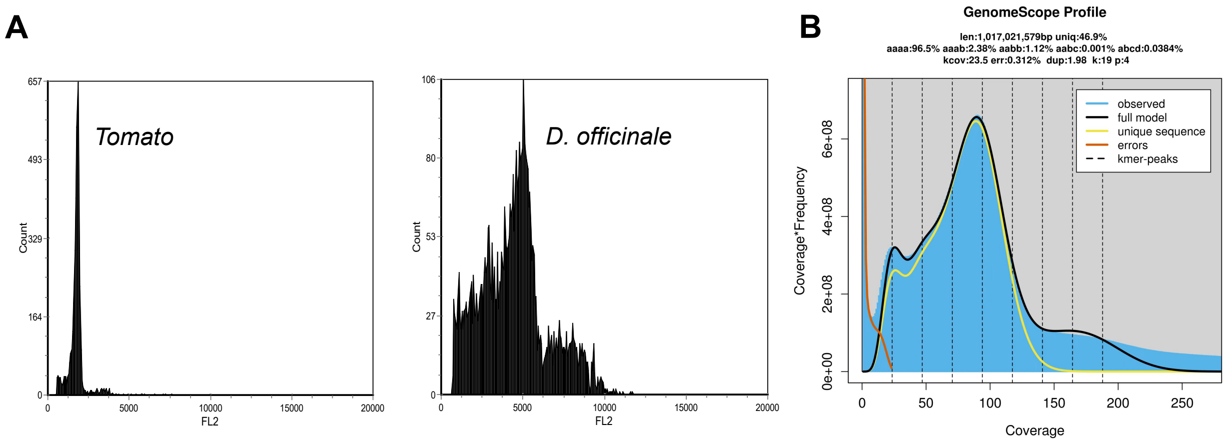


**Supplementary Figure 1.** The genome size estimation of *D. officinale*. (A) The reference species *Solanum lycopersicum* (tomato), with a nuclear DNA content of 1,654.80 Mbp, shows a fluorescence peak at 1,791.03 in the flow cytometry analysis. The *D. officinale* sample exhibits a fluorescence peak at 4,766.88, approximately 2.66 times higher than that of tomato. Based on this ratio, the estimated nuclear DNA content of *D. officinale* is approximately 4,404.30 Mbp. (B) Distribution of k-mer frequencies. The plot is generated using GenomeScope (k=19, ploidy=4).

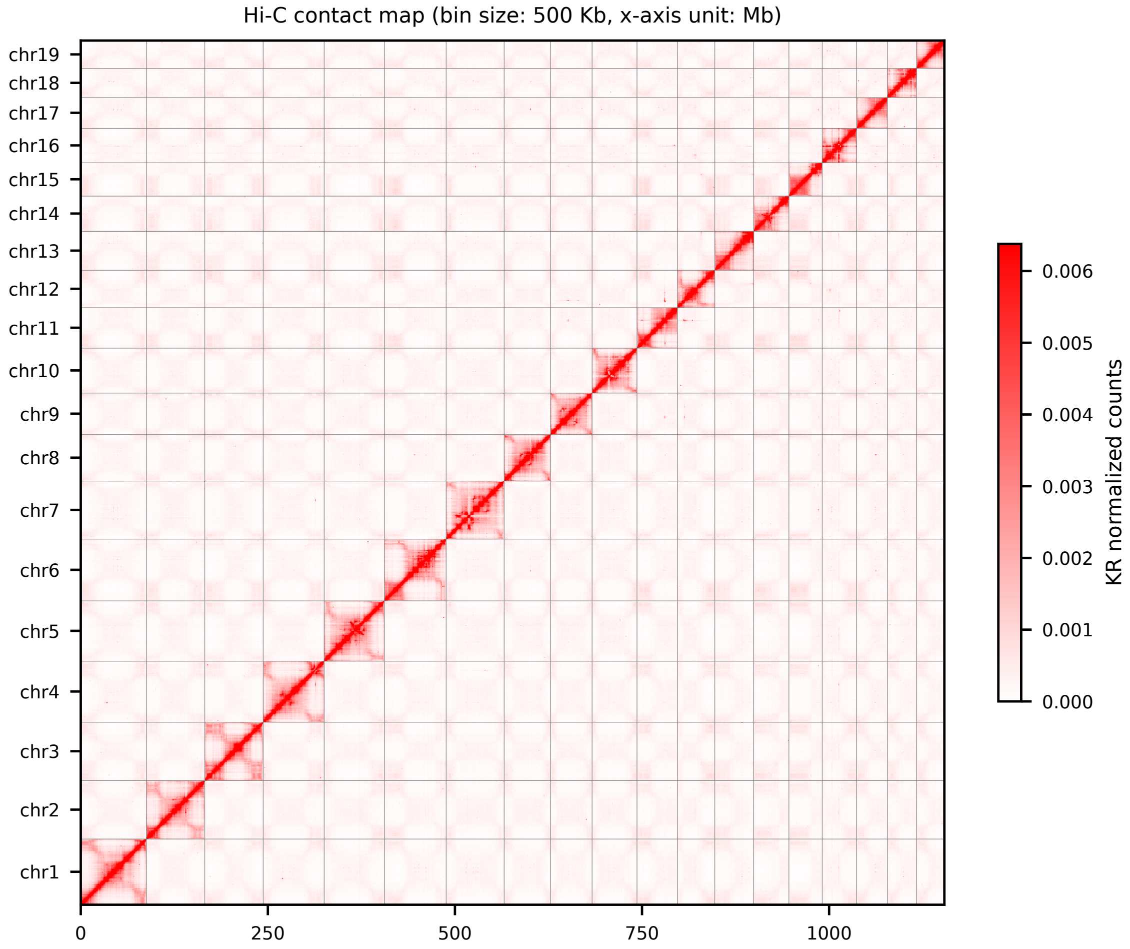


**Supplementary Figure 2.** Hi-C interaction heatmap. Hi-C interaction matrix generated at 500 kb resolution and normalized using the Knight–Ruiz (KR) method.

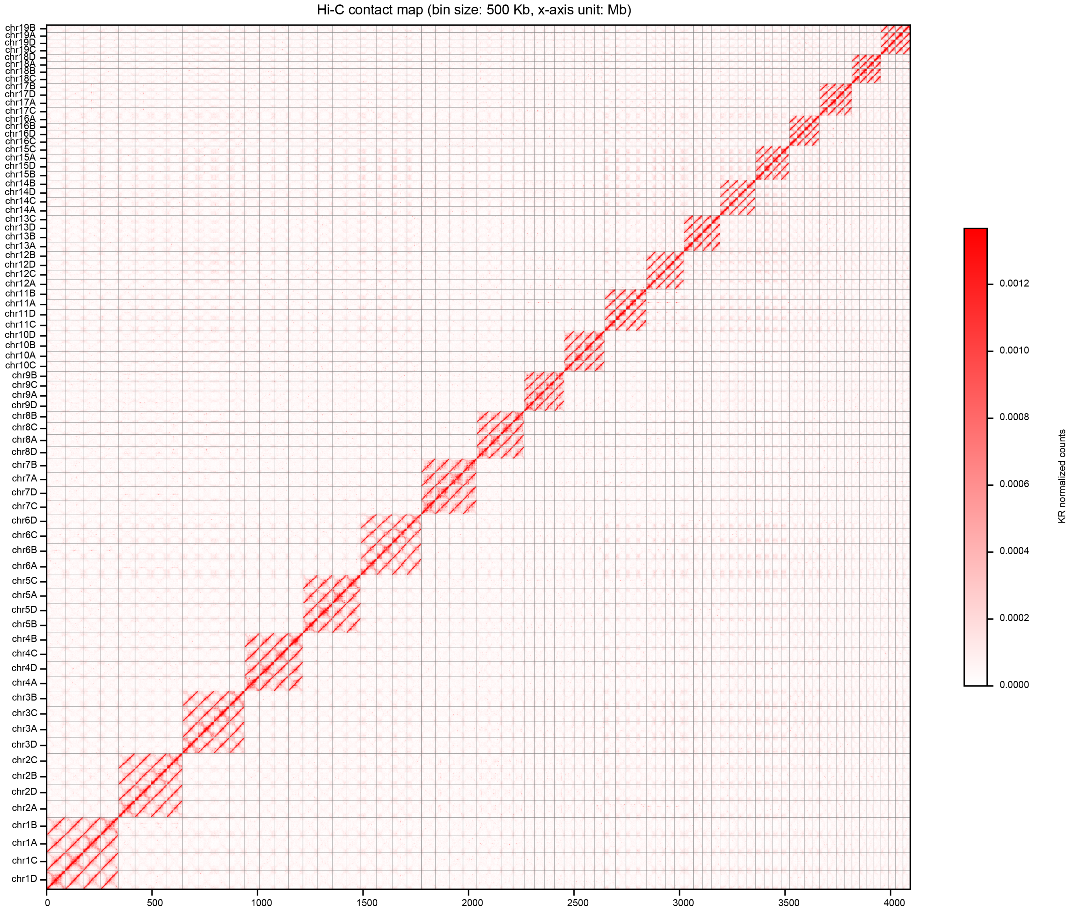


**Supplementary Figure 3.** Hi-C interaction heatmap of the haplotype-resolved genome.The Hi-C interaction matrix was generated at a 500 kb resolution and normalized using the Knight–Ruiz (KR) method. Each allelic group consists of four haplotypic chromosomes.

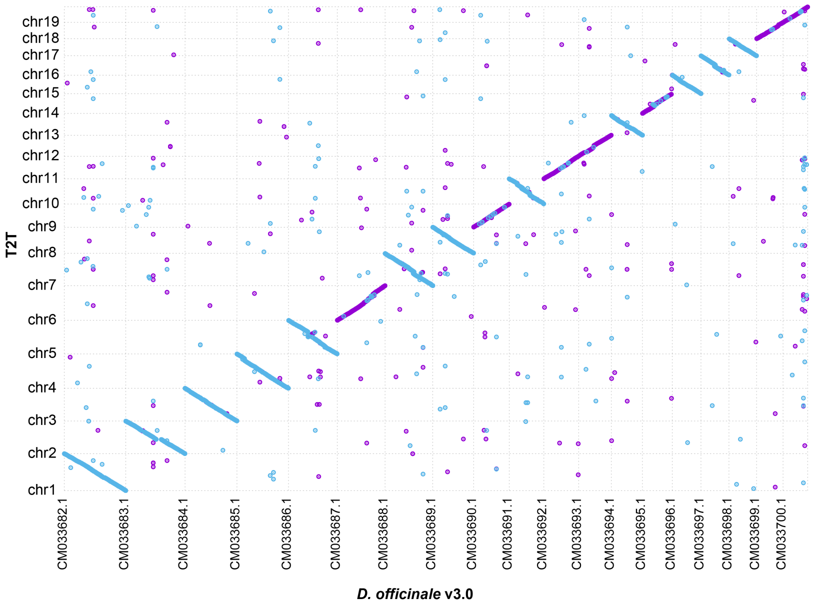


**Supplementary Figure 4.** Genome-wide MUMmer dot plot comparing *D. officinale* v3.0 and the T2T genome assembly. Each dot corresponds to a sequence alignment. Blue and purple dots denote forward and reverse alignments, respectively. *D. officinale* v3.0 sequences correspond to its 19 chromosomes.

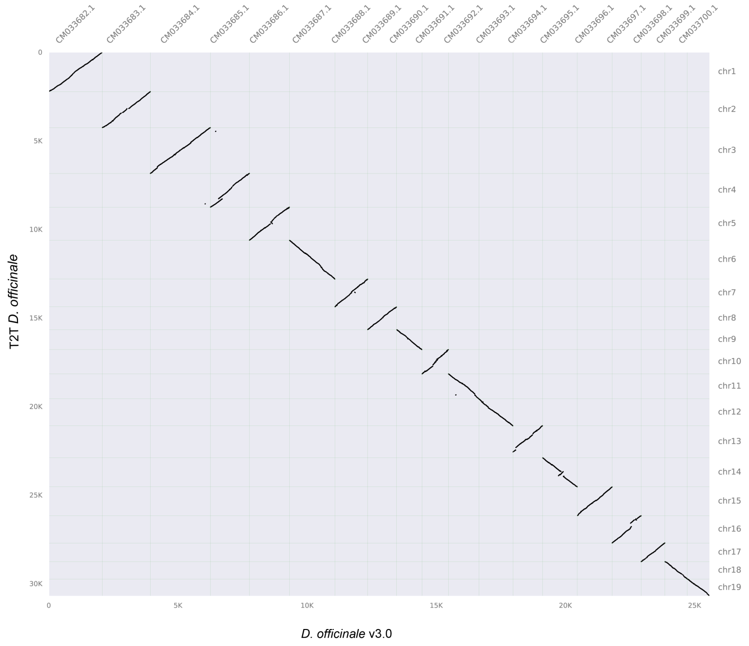


**Supplementary Figure 5.** Inter-genomic comparison between *D. officinale* v3.0 and the T2T *D. officinale* assembly was performed using JCVI. The *D. officinale* v3.0 sequences used correspond to its 19 chromosomes.

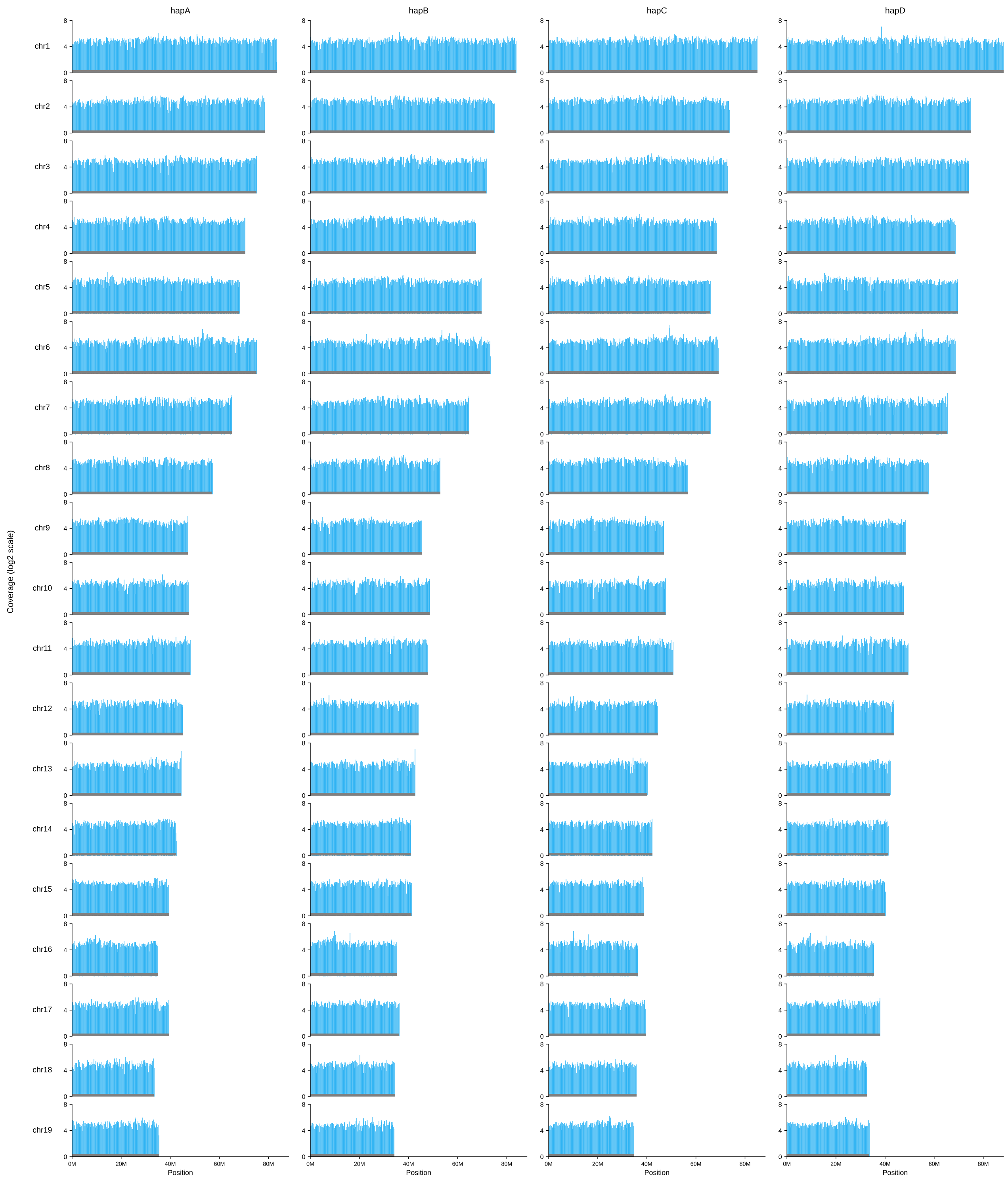


**Supplementary Figure 6.** Read coverage of PacBio HiFi reads on the *D. officinale* haplotype genome assembly, with the y-axis representing $\log_2$-mapping depth.

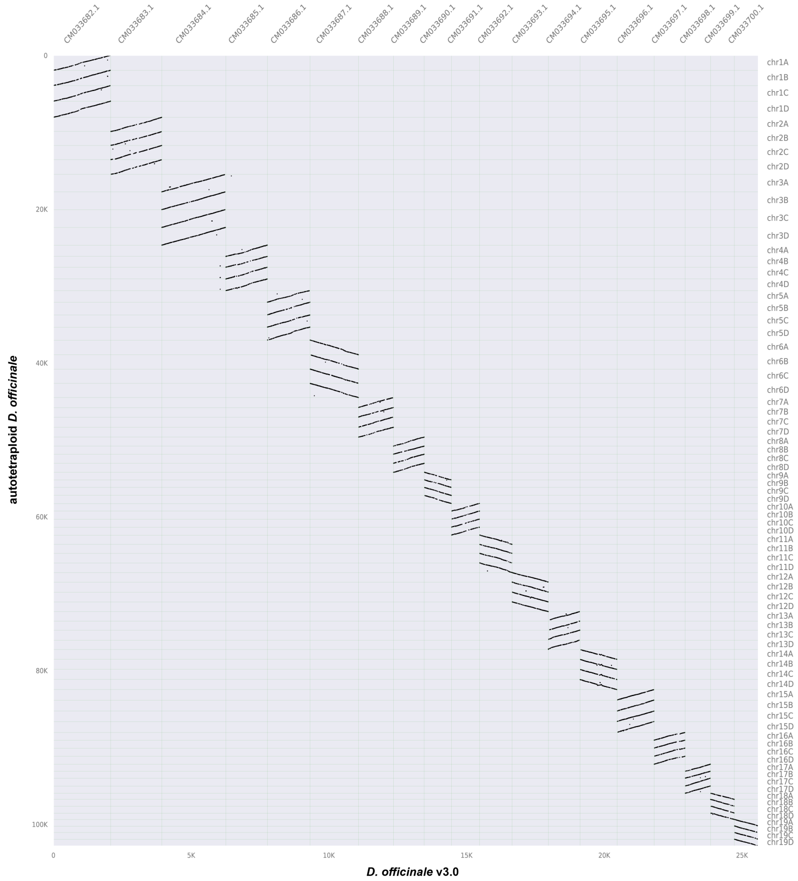


**Supplementary Figure 7.** Inter-genomic comparisons were performed between *D. officinale* v3.0 and the four haplotype assemblies using JCVI. The *D. officinale* v3.0 CDS sequences used in this analysis correspond to its 19 assembled chromosomes.

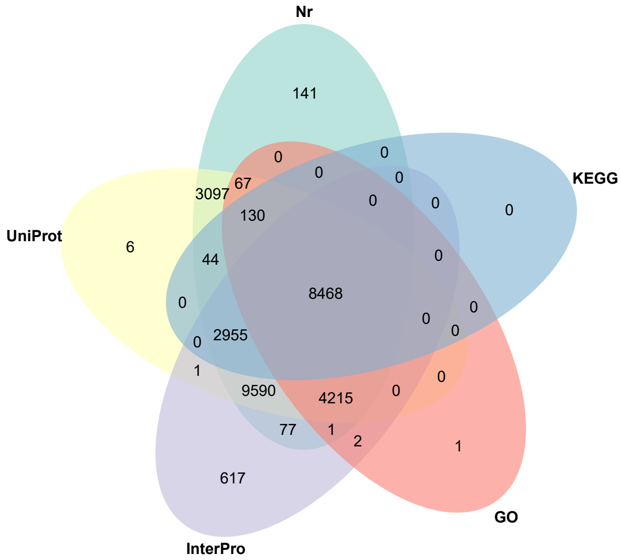


**Supplementary Figure 8.** Functional annotation of predicted genes for T2T *D. officinale* genome. Gene function annotation was performed based on five public databases: Nr, GO, KEGG, UniProt, and InterPro.

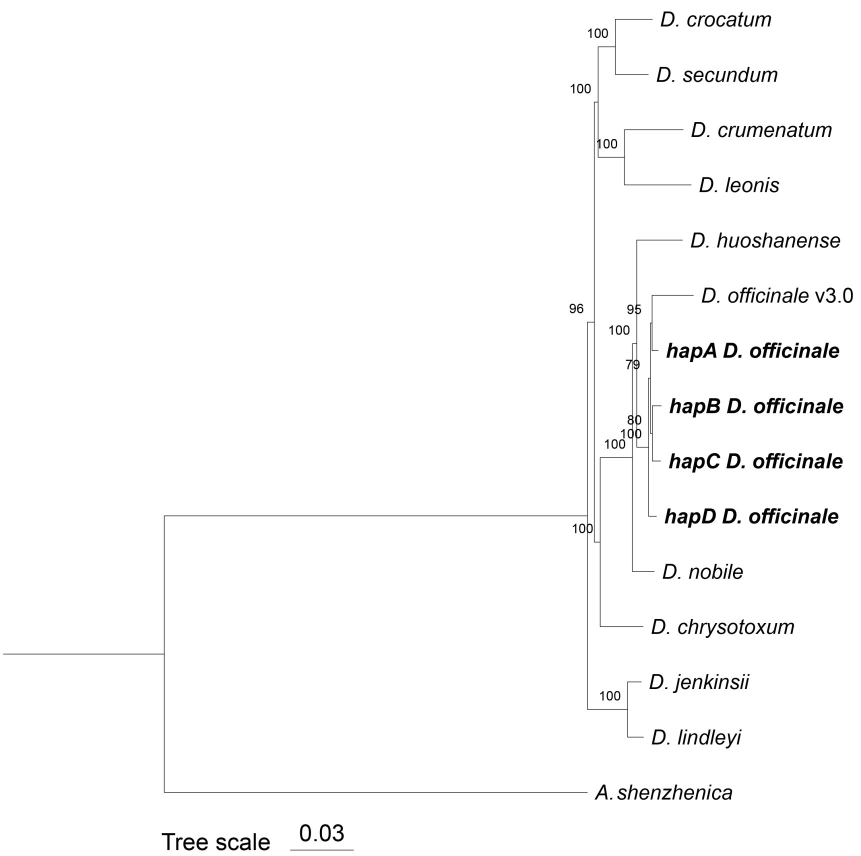


**Supplementary Figure 9.** The phylogenetic tree constructed using single-copy orthologous genes illustrates the evolutionary relationships among the four haplotypes of *D. officinale*, which were treated as independent lineages, together with other plant species. Branch support values are labeled at each node, and the published diploid *D. officinale* v3.0 genome is included.

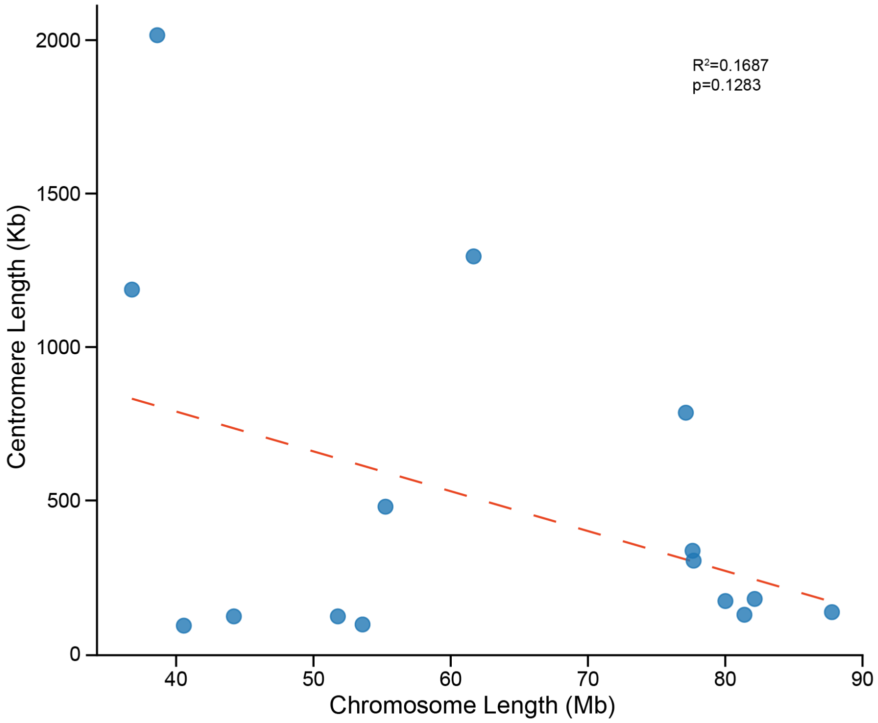


**Supplementary Figure 10.** Relationship between centromere length and chromosome length. Each point represents an individual chromosome, plotted by its total length (x-axis) against the length of its centromere (y-axis). The scatter plot illustrates the overall relationship between chromosome length and centromere length.

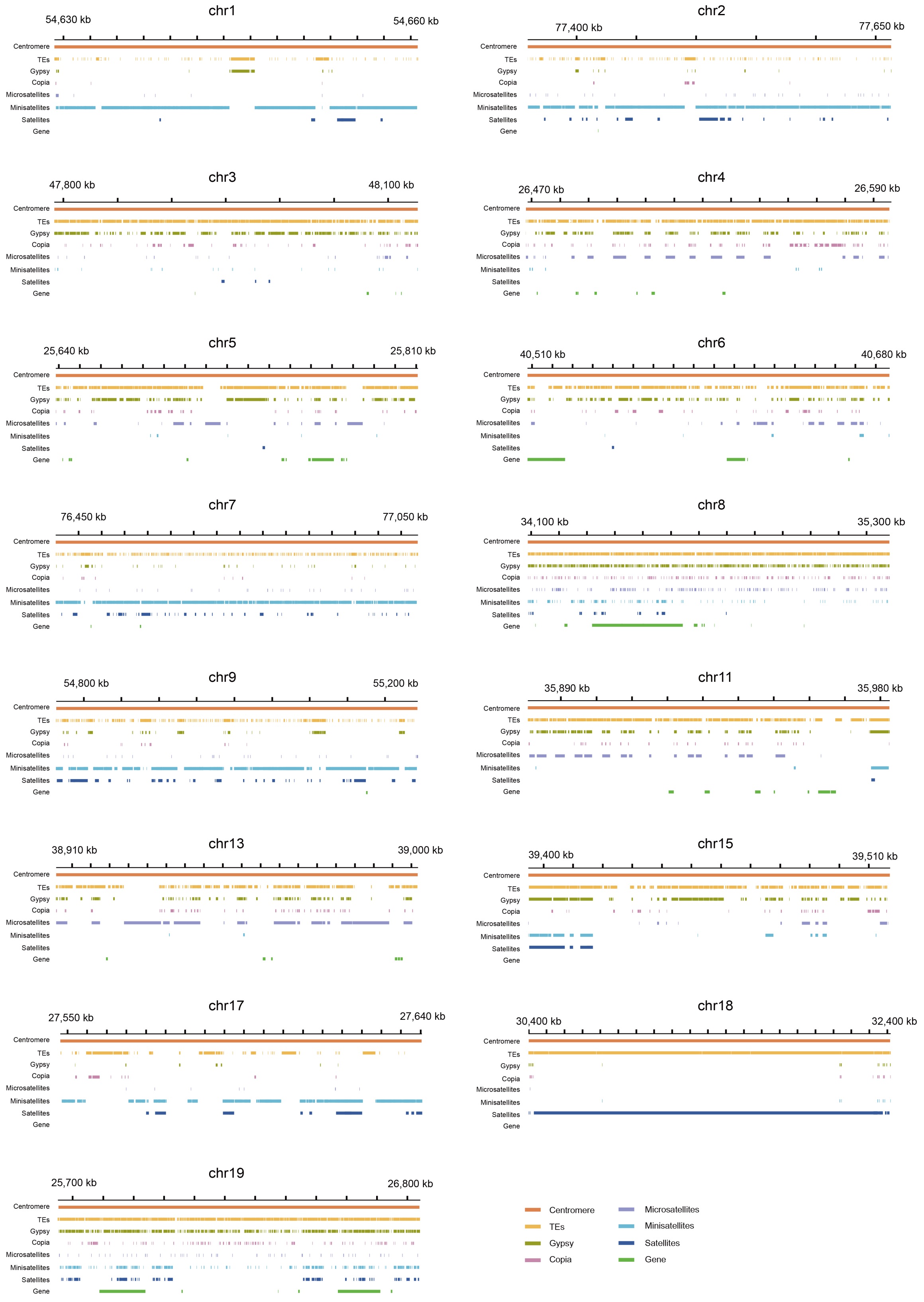


**Supplementary Figure 11.** Characterization of centromeres on the 15 chromosomes. The centromeric regions of each chromosome display the distribution of repetitive sequence types and genes. Each repeat type and the genes are displayed in separate tracks.

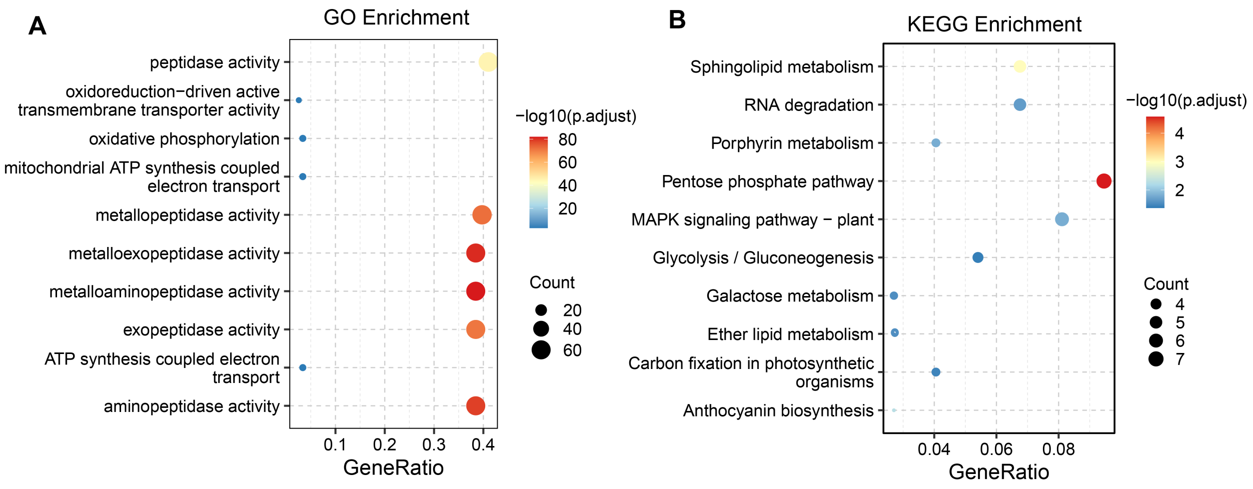


**Supplementary Figure 12.** (A) GO and (B) KEGG enrichment analysis of genes in the species-specific gene families of *D. officinale*

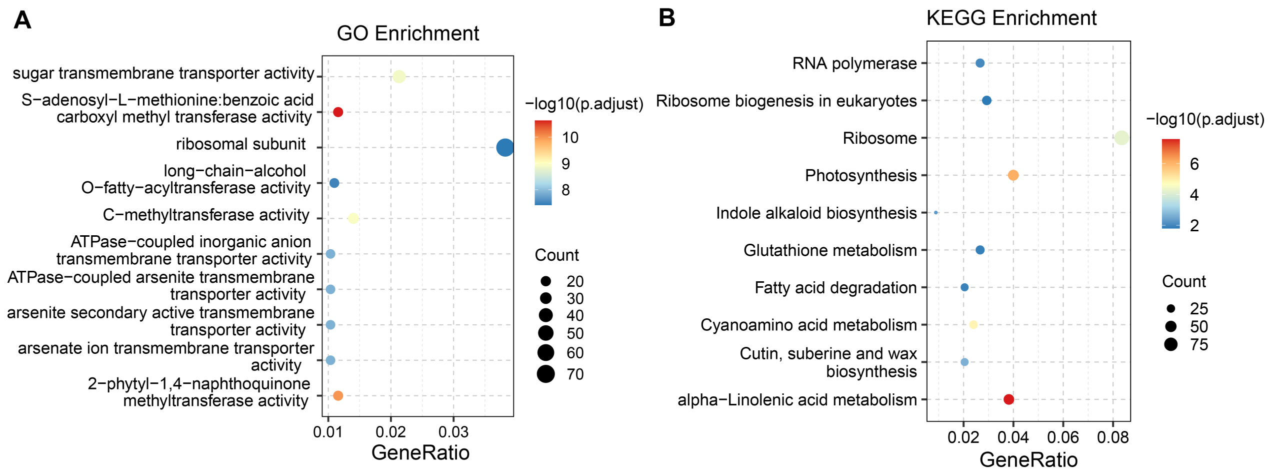


**Supplementary Figure 13.** (A) GO and (B) KEGG enrichment analysis of genes in the expanded gene families.

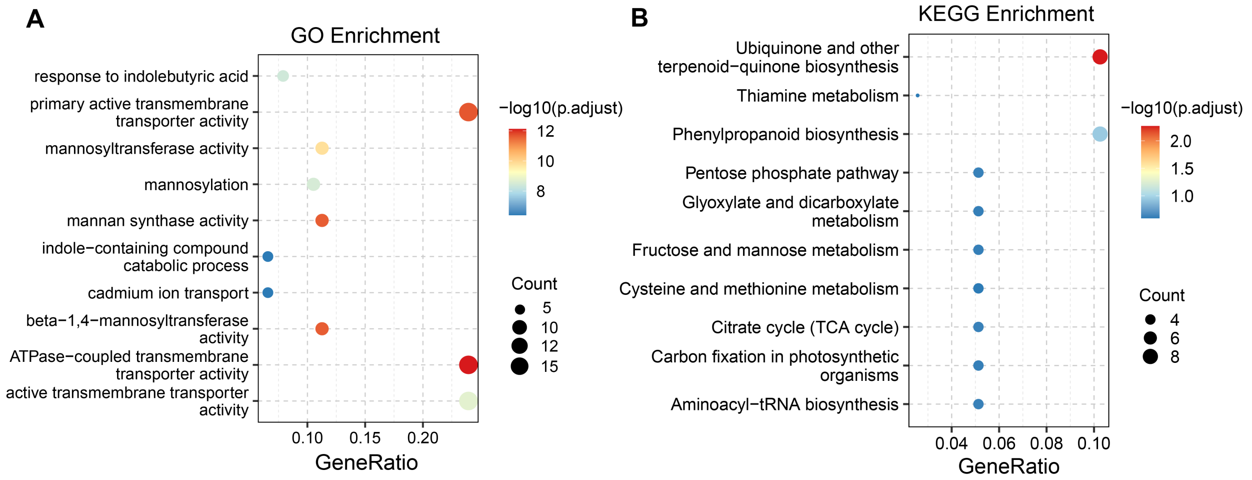


**Supplementary Figure 14.** (A) GO and (B) KEGG enrichment analysis of genes in the contracted gene families.

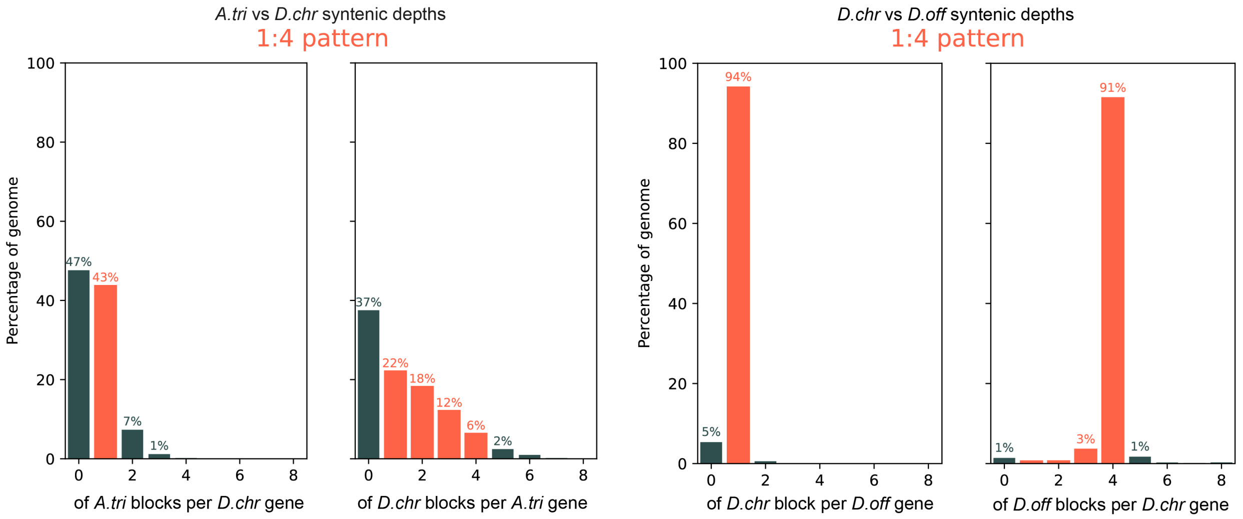


**Supplementary Figure 15.** Syntenic depth patterns based on JCVI analysis. A 1:4 syntenic depth pattern is observed between *A. trichopoda* and *D. chrysotoxum* (left), and between *D. chrysotoxum* and tetraploid *D. officinale* (right).

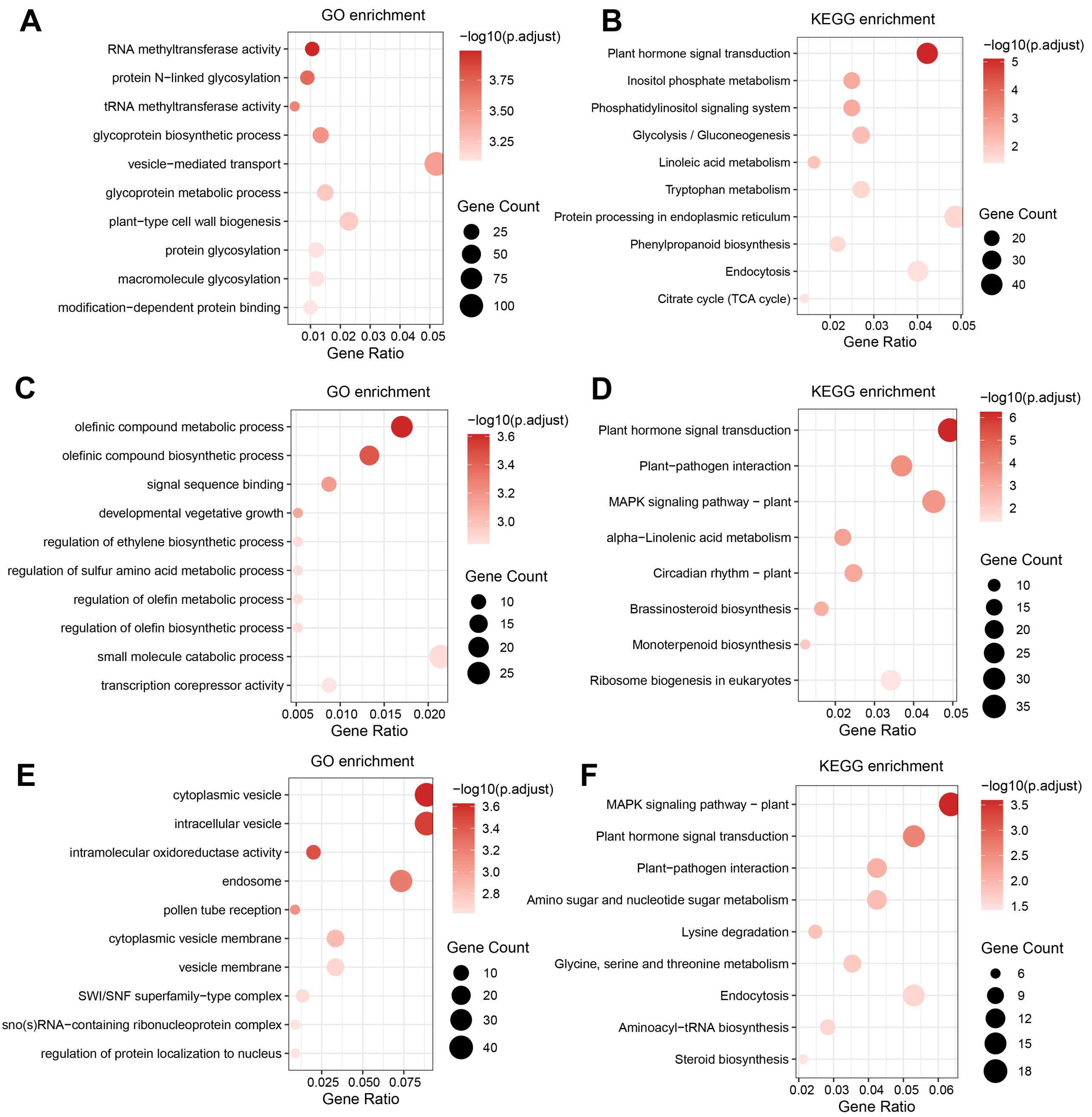


**Supplementary Figure 16.** Functional enrichment analysis of different allelic expression patterns in root tissue. (A–B) GO and KEGG enrichment analyses of genes showing dual-allele dominant expression. (C–D) GO and KEGG enrichment analyses of genes showing single-allele suppressed expression. (E–F) GO and KEGG enrichment analyses of genes showing balanced expression. All analyses were performed using the hapA gene copies corresponding to each allelic expression category. The Y-axis lists the enriched functional terms or pathways. The x-axis represents the Gene Ratio

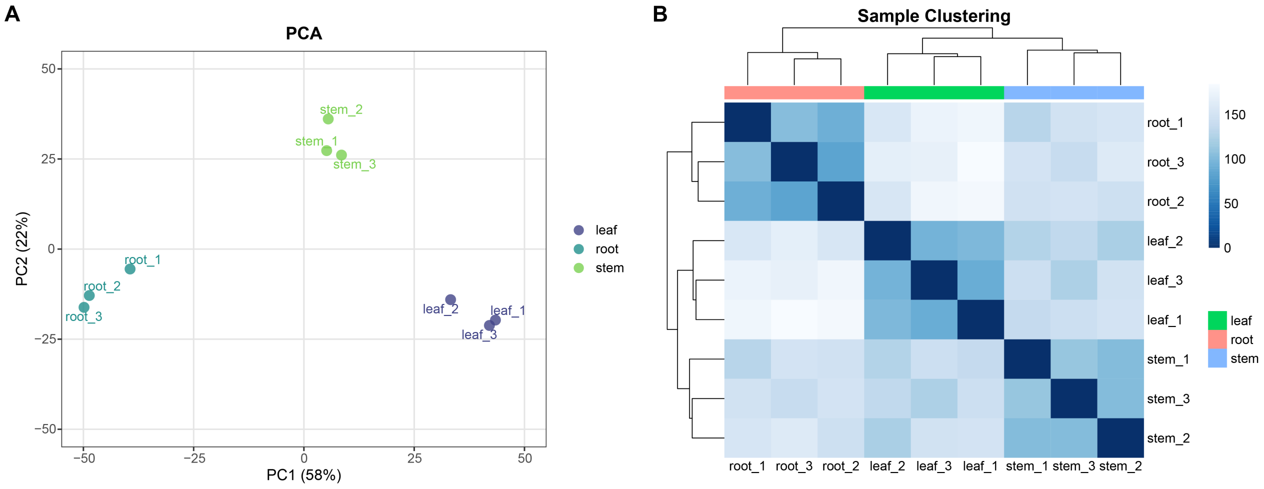


**Supplementary Figure 17.** principal component analysis (PCA) and Sample clustering based on transcriptome profiles. (A) PCA of all RNA-seq samples based on variance-stabilizing transformation (VST) of gene expression values. (B) Hierarchical clustering heatmap of samples based on pairwise Euclidean distances computed from VST-transformed expression data. The color gradient represents the degree of dissimilarity between samples

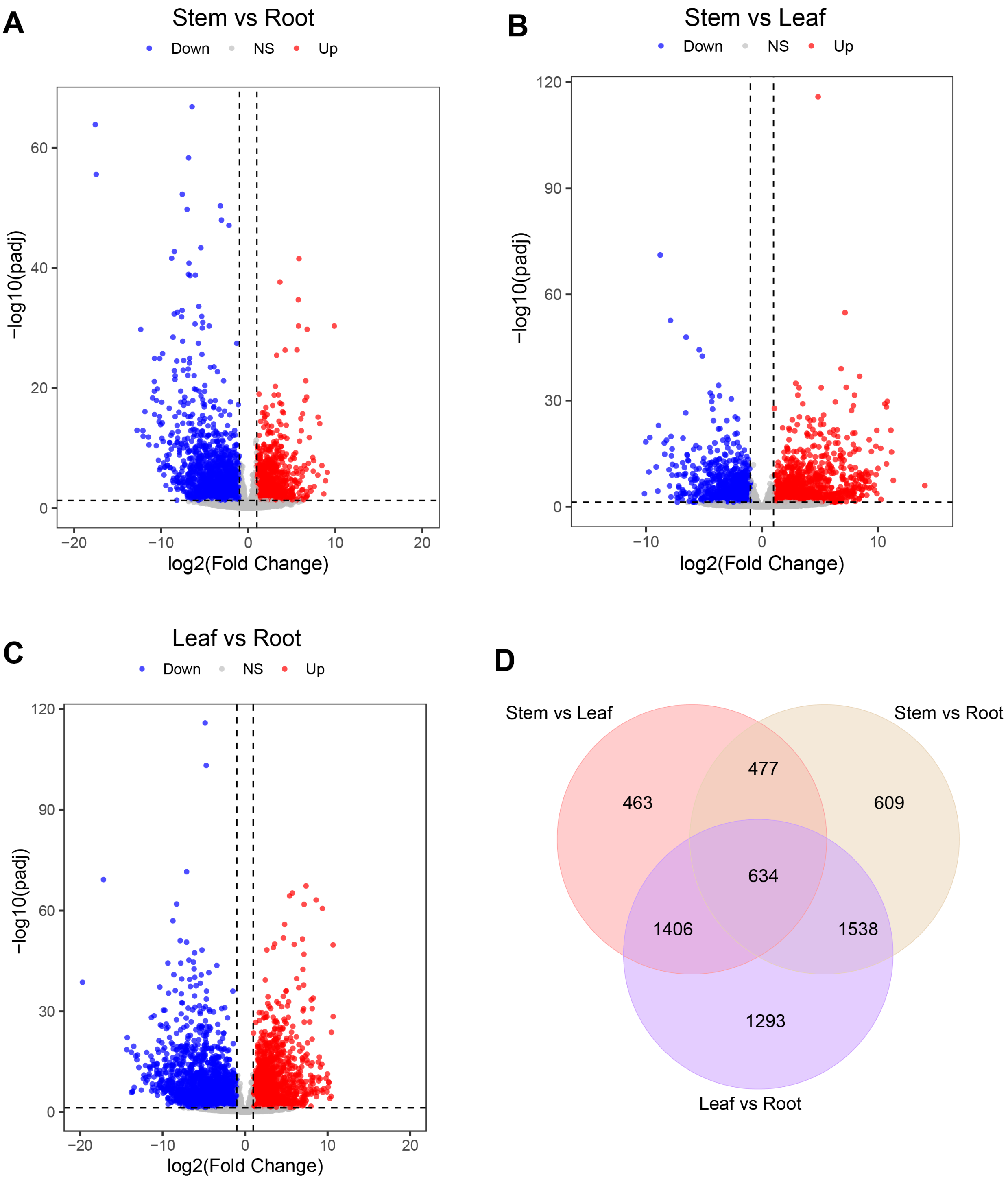


**Supplementary Figure 18.** Differentially expressed genes (DEGs) in three tissues of *D. officinale*. (A) Volcano plot showing differential expression analysis between stem and root. (B) Volcano plot showing differential expression analysis between stem and leaf. (C) Volcano plot showing differential expression analysis between root and leaf. Red and blue dots indicate significantly up-regulated and down-regulated genes, respectively. Grey dots indicate non-significant genes. (D) Venn diagram showing the overlap of DEGs identified in the three pairwise comparisons ($|\log_2 \mathrm{FC}| > 1$, $p_{\mathrm{adj}} < 0.05$).

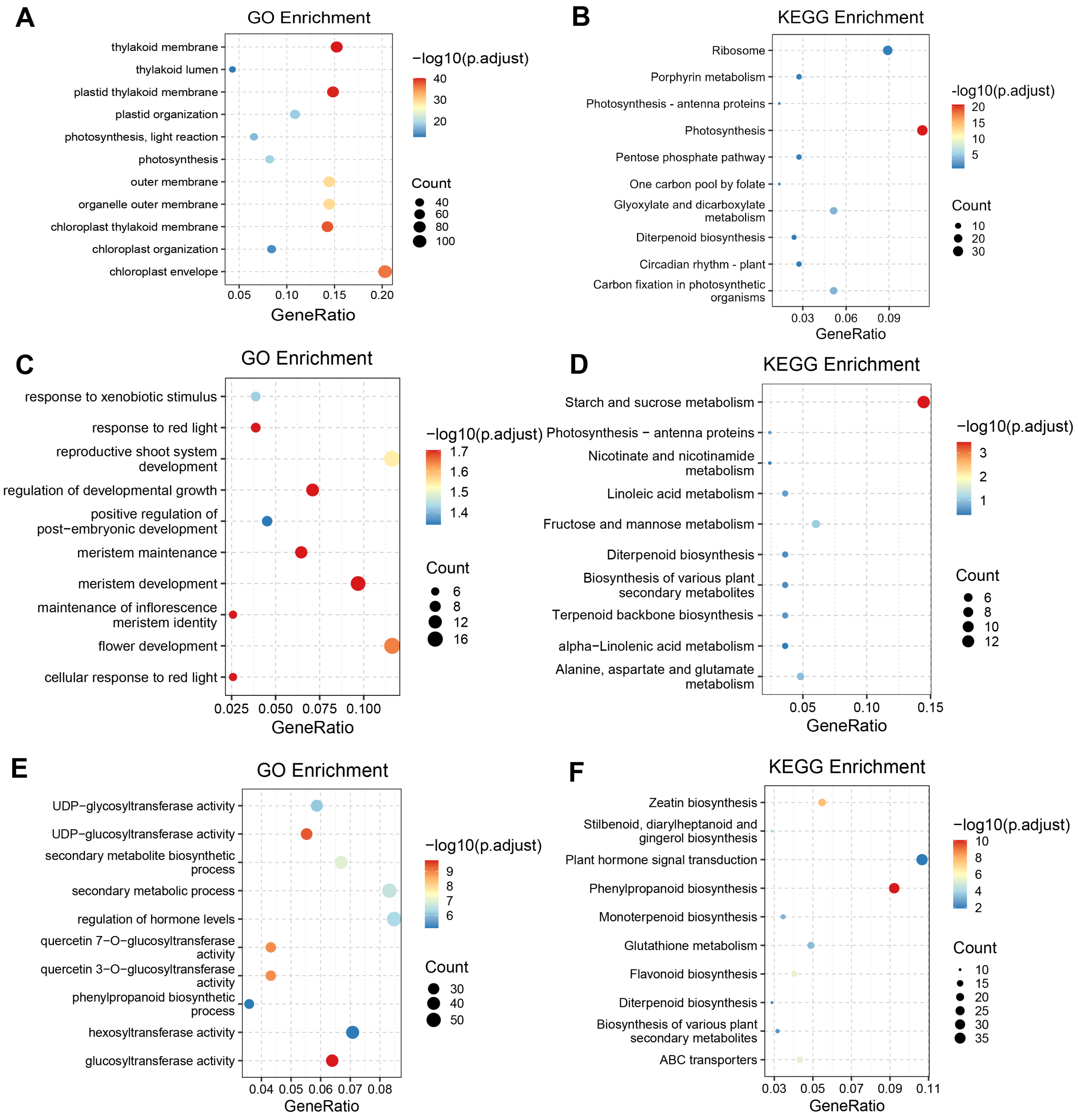


**Supplementary Figure 19.** GO and KEGG enrichment analyses of genes up-regulated in three tissues of *D. officinale*. (A, B) GO (A) and KEGG (B) enrichment analyses of genes up-regulated in leaf relative to stem and root. (C, D) GO (C) and KEGG (D) enrichment analyses of genes up-regulated in stem relative to leaf and root. (E, F) GO (E) and KEGG (F) enrichment analyses of genes up-regulated in root relative to leaf and stem. The x-axis represents the gene ratio, the size of each dot indicates the number of genes enriched in each term, and the color gradient indicates the significance level expressed as $-\log_{10}$(adjusted $p$-value).

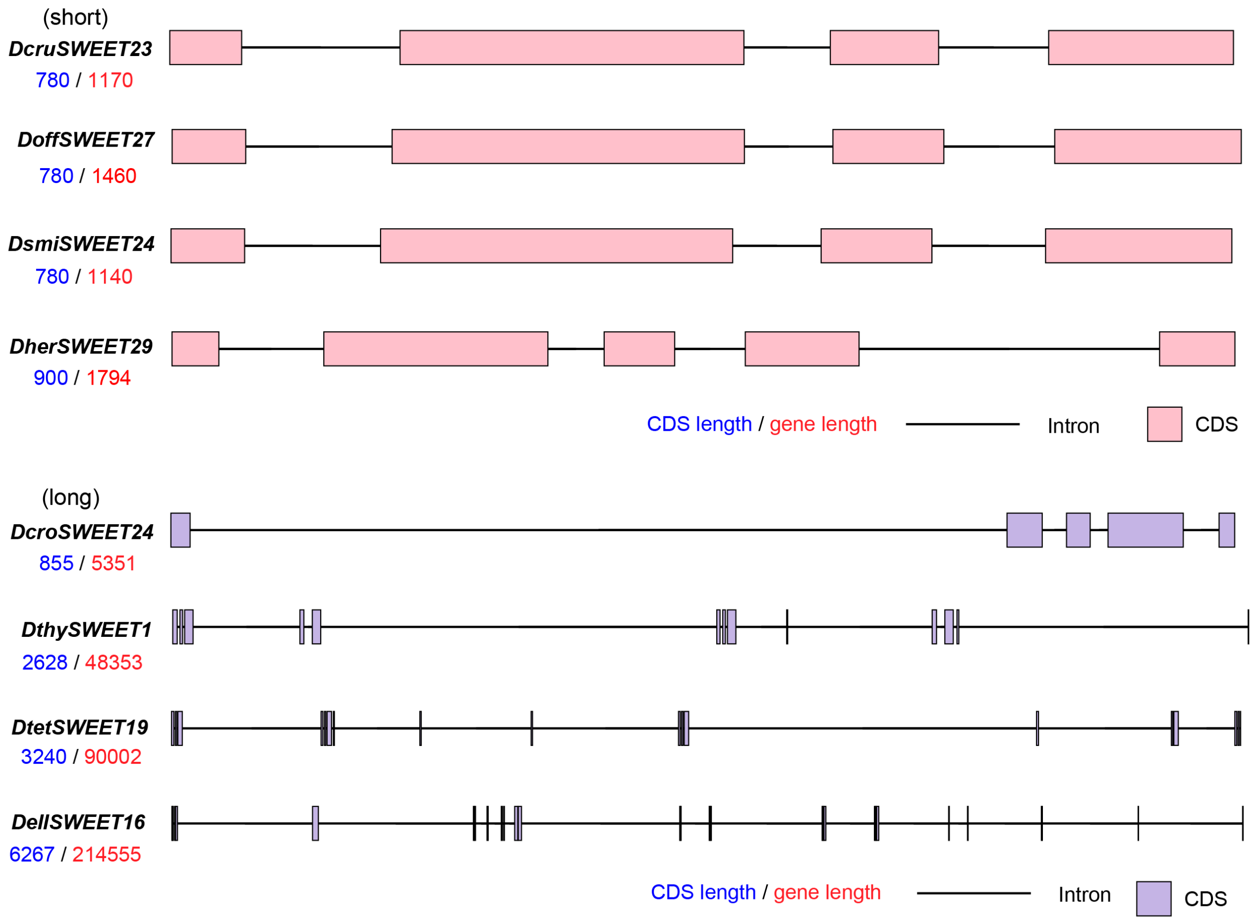


**Supplementary Figure 20.** CDS–intron structure of *SWEET* genes. Rectangles represent CDSs, and black lines represent introns. Blue labels indicate CDS length, and red labels indicate total gene length. Four relatively short genes (*DcruSWEET23*, *DoffSWEET27*, *DsmiSWEET24*, and *DherSWEET29*) and four relatively long genes (*DcroSWEET24*, *DthySWEET1*, *DtetSWEET19*, and *DellSWEET16*) were selected for comparison. To clearly illustrate gene structure organization, all genes are schematically illustrated at a uniform length and are not drawn to scale.

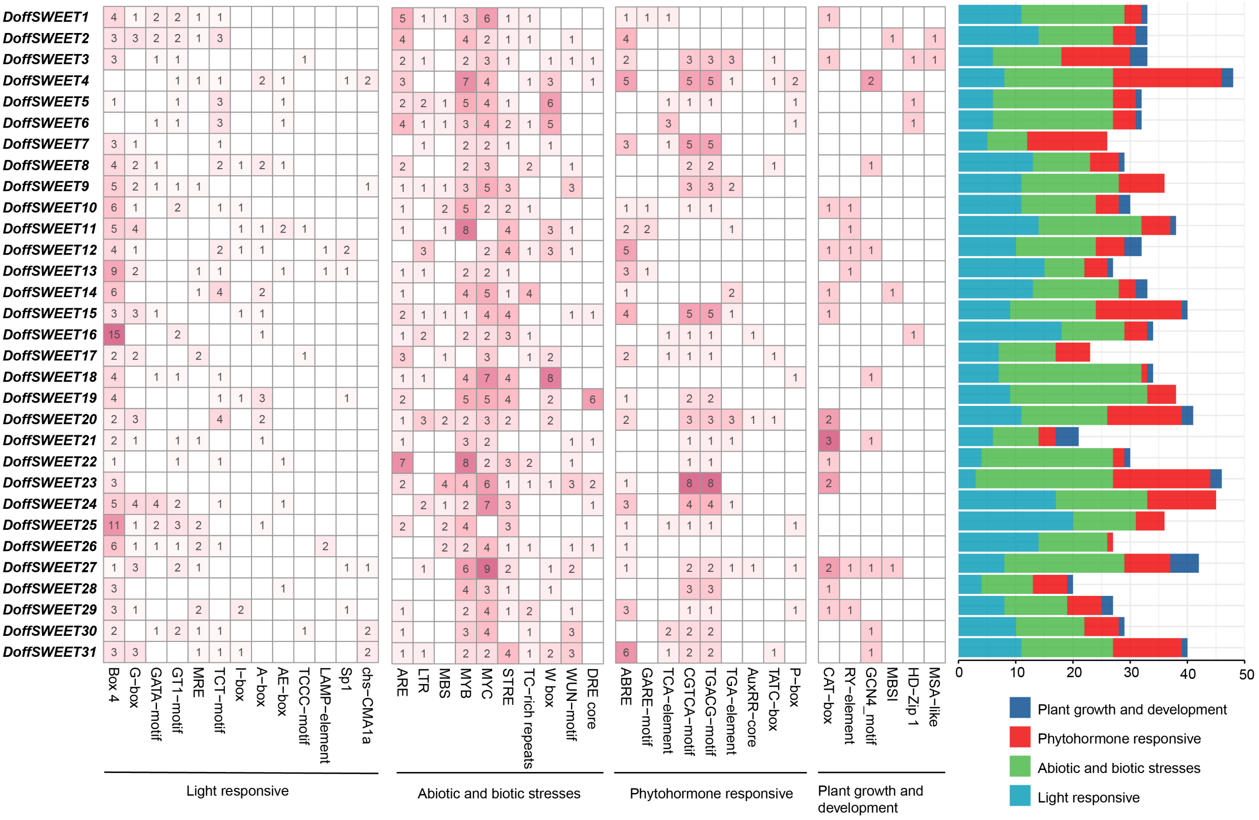


**Supplementary Figure 21.** Cis-acting elements identified in the promoter regions of *DoffSWEET* genes in *D. officinale*. Heatmap showing the counts of individual cis-elements for each gene, with the number in each cell indicating the occurrence of the corresponding element. Stacked bar chart summarizing the total counts of cis-elements in four functional categories: light responsive (blue), abiotic and biotic stresses (green), phytohormone responsive (red), and plant growth and development (cyan).
